# Genetic requirements for repair of lesions caused by single genomic ribonucleotides in S phase

**DOI:** 10.1101/2022.06.30.498227

**Authors:** Natalie Schindler, Matthias Tonn, Vanessa Kellner, Jia Jun Fung, Arianna Lockhart, Olga Vydzhak, Thomas Juretschke, Stefanie Möckel, Petra Beli, Anton Khmelinskii, Brian Luke

**Affiliations:** Johannes Gutenberg University Mainz, Institute for Developmental Neurology (IDN), Biozentrum 1, Hanns-Dieter-Hüsch-Weg 15, 55128 Mainz, Germany; Institute of Molecular Biology (IMB), Ackermannweg 4, 55128 Mainz, Germany; Department of Biology, New York University, New York, NY, USA

**Author notes:** these authors contributed equally to the manuscript (Natalie Schindler, Matthias Tonn). Co-authors e-mail addresses.

## Abstract

Single ribonucleoside monophosphates (rNMPs) are transiently present in eukaryotic genomes. The RNase H2-dependent ribonucleotide excision repair (RER) pathway ensures error-free genomic rNMP removal. In pathological conditions, genomic rNMP levels can rise and persist. If these rNMPs hydrolyse in, or prior to, S phase, toxic single-ended double-strand breaks (seDSBs) can occur upon an encounter with replication forks. How such rNMP-derived seDSB lesions are repaired is unclear. We employed a cell cycle phase restricted allele of RNase H2 as a genetic tool to induce nicks at rNMPs specifically in S phase to generate such lesions and study their repair. Here, we introduce a network of genes that maintain DNA integrity when rNMP-derived nick lesions arise during DNA replication. We use genetic methods to characterise the molecular requirements of a Top1-independent, rNMP-derived <u>n</u>ick lesion repair (NLR). In NLR, the *RAD52* epistasis group becomes essential for homology-directed repair (HDR). Moreover, the previously described Rtt101^Mms1-Mms22^ dependent ubiquitylation of histone H3 is essential for NLR in cells with high rNMP load, and loss of Rtt101^Mms1-Mms22^ combined with RNase H2 dysfunction leads to compromised cellular fitness. We discuss the genetic NLR network in the context of human disease, where cancer therapies may be able to exploit these synthetic lethalities.

## Introduction

Single ribonucleoside monophosphates (rNMPs) are present in the genomic DNA (gDNA) of all organisms. The budding yeast *Saccharomyces cerevisiae* incorporates about 10,000 rNMPs into the genome per cell cycle (Nick McElhinny et al., 2010). The DNA polymerases transiently incorporate rNMPs during DNA replication. The double-stranded context of gDNA does not allow the reduction of the 2’-hydroxyl (2’-OH) group in the misincorporated rNMP to reduce it to a deoxy group, hence its presence can lead to single-strand DNA (ssDNA) breaks (nicks) that result in replication fork collapse and the formation of single-ended double-strand breaks (seDSBs) in S phase. In addition, genomic rNMPs themselves hinder the passage of DNA polymerases and cause replication stress from yeast to human cells (Hiller et al., 2012; Lazzaro et al., 2012; Nick McElhinny et al., 2010; Pizzi et al., 2015; Williams et al., 2013; Zimmermann et al., 2018). Therefore, it is critical to remove rNMPs in a timely manner to prevent rNMP-derived genomic instability.

RNase H2 is the central ribonucleotide excision repair (RER) enzyme in yeast and mammalian cells (Cerritelli & Crouch, 2009). The majority of rNMPs that have been incorporated into gDNA during DNA replication in S phase are removed in the subsequent G2 phase (Lockhart et al., 2019). When RER fails, topoisomerase 1 (Top1) can process genomic rNMPs. However, this activity is associated with genomic instability due to error-prone branching in the Top1 pathway (Kim et al., 2011; Sekiguchi & Shuman, 1997). Even in the presence of RER, Top1 nicks some rNMPs (Reijns et al., 2022), but the precise interplay between RNase H2 and Top1 remains to be elucidated. Nonetheless, timely elimination of genomic rNMPs is crucial, and defective RER is associated with human diseases such as chronic lymphocytic leukemia and prostate cancer (Crow et al., 2006; Zimmermann et al., 2019).

RNase H2 is a trimeric enzyme in yeast (*RNH201, RNH202, RNH203*) and mammalian cells (RNASEH2A, RNASEH2B, RNASEH2C) (Cerritelli & Crouch, 2009). We previously engineered cell cycle regulated alleles for the *RNH202* gene to restrict expression of the enzyme to either the S or G2 phase of the cell cycle (Lockhart et al., 2019). In addition to the finding that the expression of *RNH202* exclusively in G2 was sufficient to suppress RER defects, we observed an unexpected fitness defect when RNase H2 activity was restricted to S phase (Lockhart et al., 2019). Yeast with S phase expressed *RNH202* (*S-RNH202-TAP, referred to as S-RNH202*) experienced toxicity caused by nicking of the gDNA and relied on the homology-directed repair (HDR) factor Rad52 for survival (Lockhart et al., 2019). The decreased fitness of the *S-RNH202* strain was strongly exacerbated in presence of the *pol2-M644G* allele, a Polymerase ε (Pol ε) mutant that incorporates 10-fold more rNMPs (Nick McElhinny et al., 2010; Williams et al., 2016). Notably, the *S-RNH202* phenotype was independent of Top1 activity suggesting that nicking of rNMPs in the S phase causes toxic seDSBs during replication.

Surprisingly, rNMP accumulation can be tolerated in the absence of both RER and Top1 pathways. This is evidenced by the viability of budding yeast lacking Top1 and expressing an allele of RNase H2 that is deficient in RER (but proficient in R-loop removal) (Chon et al., 2013). This implies that cells can tolerate the presence of replication stress and DNA damage from rNMPs, or that there might be another rNMP lesion repair pathway that is independent of RNase H2 and Top1 (discussed in (Kellner & Luke, 2020)).

In this study, we set out to get a better understanding of how rNMP-induced DNA lesions are repaired, when the nicking occurs during, or prior to, DNA replication. We used the *S-RNH202* allele as a molecular tool to promote nicking of genomic rNMPs in S phase (Lockhart et al., 2019). Using synthetic genetic array (SGA) technology (Tong et al., 2001), we demonstrate that the *RAD52* HDR epistasis group, the histone remodelers genes *Asf1* and *Rtt109*, the STR (Sgs1-Top3-Rmi) complex, the Mus81-Mms4 resolvase, and the E3-Ubiquitin ligase complex *Rtt101*, *Mms1,* and *MmsS22* are all required for the tolerance of rNMP-derived nicks during S phase. These factors comprise a Top1-independent, rNMP-derived <u>n</u>ick lesion repair (NLR) pathway. We also found that histone H3 ubiquitylation by the replisome-associated Rtt101^Mms1-Mms22^ complex is critical for NLR in high rNMP conditions, pointing to a role for chromatin remodeling in NLR.

We summarize our genetic data in a descriptive model that represents our idea of the molecular processes in the NLR repair pathway. When a replication fork runs into an rNMP-derived leading strand nick, a seDSB is formed. Locally, Rtt101-dependent post-translational modifications at chromatin and elsewhere then take place that may support resection of the strand end in preparation for HDR. The RST and Mus81-Mms4 complexes then provide resolution of the recombination intermediates. In the course of the NLR pathway, the rNMP that initiated the strand breakage was removed, making NLR a *bona fide* rNMP repair. Importantly, we report a negative genetic interaction between *RTT101* and RNase H2, which becomes synthetic lethal when the genomic rNMP load increases. These data in yeast may provide therapeutic insights and alternatives for human cancer treatment in genetic contexts where RNase H2 is dysfunctional such as RER-deficient cancers (Zimmermann et al., 2018).

## Results

### Synthetic lethal screen identifies a network required for rNMP-derived lesion tolerance in S phase

We employ the *S-RNH202-TAP* allele (from here on referred as *S-RNH202*) as a genome-wide tool to endogenously nick genomic rNMPs in S phase. Restricting the expression of RNase H2 to S phase also results in the accumulation of genomic rNMPs as canonical RER occurs outside of the S phase, hence the rNMP load is similar in *S-RNH202* and in RER-deficient strains as measured by alkaline gel electrophoresis (Lockhart et al., 2019). The *S-RNH202* allele also presents the same rate of mutagenesis as the RNase H2 deletion (*rnh202*Δ) in the presence of the *pol2-M644G* allele, a Polymerase ε (Pol ε) mutant that increases the rNMP load by 10-fold (Williams et al., 2016) (***Figure S1A***). RNase H2 deletion and *S-RNH202* expressing strains share not only the same amount of genomic rNMPs, and the same mutagenesis rate but also are both highly sensitive towards hydroxyurea (HU) (in the *pol2-M644G* background) (***Figure S1B***). We have shown before that methyl methane sulfonate (MMS) stabilizes R-loops which are potentially toxic RNA-DNA hybrids that are removed by RNase H1 and RNase H2 (Lockhart et al., 2019). In the presence of MMS, the *rnh1*Δ *rnh201*Δ double mutant and *rnh1*Δ *S-RNH202* double mutant are inviable (***Figure S1C***). Therefore, both canonical RNase H2 functions, R-loop removal and RER, occur outside of S phase. Hence, employing the *S-RNH202* allele as an enzymatic tool to endogenously nick genomic rNMPs recapitulates many phenotypes of an RNase H2 deletion, and suggests that many problems associated with loss of RER are due to rNMP nicking during DNA replication. Therefore, the *S-RNH202* allele is relevant both in terms of understanding rNMP repair during RER deficiency (rNMP hydrolysis in S phase) and in canonical RER, when RNase H2 nicked rNMPs are not repaired in a timely manner and are encountered in the following S phase.

To identify factors involved in repair of rNMP-derived lesions occurring in S phase we performed a synthetic genetic array (SGA) analysis cell cycle restricted alleles of *RNH202* in the budding yeast *Saccharomyces cerevisiae* (***Figure 1A***). We generated G1-, S-, and G2-restricted alleles of *RNH202* in the query background (***Figure S1D, S1E***). Then, we crossed the three queries and the wild type control to the haploid yeast knockout collection (YKO) of all non-essential yeast genes. We derived haploid double mutants from the resulting diploid strain and determined their fitness by measuring colony size (***Figure 1A***). We compared the hits of each *RNH202* cell cycle allele with the wild type *RNH202* control to identify allele-specific genetic interactions (representative examples ***Figure S1F-S1H***). Out of 4790 gene knockouts included in the screen, we identified 21 synthetic sick interactions for the *G1-RNH202* allele (***Figure 1B***), 45 for the *S-RNH202* allele (***Figure 1C***), and eight for the *G2-RNH202* allele (***Figure 1D***). Of those hits, five genes were essential to support normal colony size among all three alleles (***Figure 1E***). Gene Ontology (GO) revealed that the GO processes related to “DNA recombination” and “DNA repair” were enriched among the 45 synthetic sick interactions of the *S-RNH202* allele (***Figure 1F***). This is in line with our previous finding that the HDR factor Rad52 is essential in the *S-RNH202* genetic background (Lockhart et al., 2019). We tested all candidates by manual tetrad dissection and curated the genetic interaction network accordingly (***Table S1, Figure 1G***). Among the synthetic sick interactions unique to the *S-RNH202* allele, we identified *RAD52* epistasis group genes (*RAD52, RAD54, RAD55, RAD57*) consistent with our previous report (Lockhart et al., 2019), the *MUS81-MMS4* nuclease complex, the *RMI1-SGS1-TOP3* (RST) helicase complex, the *MRE11-XRS2-RAD50* (MRX) nuclease complex, the nucleosome assembly factors *RTT109 and ASF1* and the *RTT101^MMS1^* ubiquitin ligase (***Figure 1G***). None of the G2-specific synthetic sick interactions was confirmed by manual tetrad dissection and thus were false-positives, consistent with canonical RER occurring in this phase of the cell cycle (see ***Figure S1I*** for examples). Surprisingly, only two hits were confirmed with the *G1-RNH202* allele, both involved in the HDR pathway. The G1 allele is the least tightly regulated of all *RNH202* alleles (***Figure S1E***). To exclude that the complementation of *G1-RNH202* was due to a weak expression of the G1 allele into S phase we performed synchronization experiments combined with induced-expression of RNase H2 only in G1 phase. These unpublished data support the spotting in ***Figure S1B*** and will be part of the future characterizing of RNase H2 activity in the G1 phase. In summary, the results with the *G1-RNH202* allele indicate that RNase H2 initiated rNMP-repair may also take place in G1 phase. However we could envisage that HDR is needed to an extent, in the case that nicked rNMPs from G1 are passed into the following S phase where these nicks again meet the replisome and ultimately would form seDSBs.

**Figure 1.**
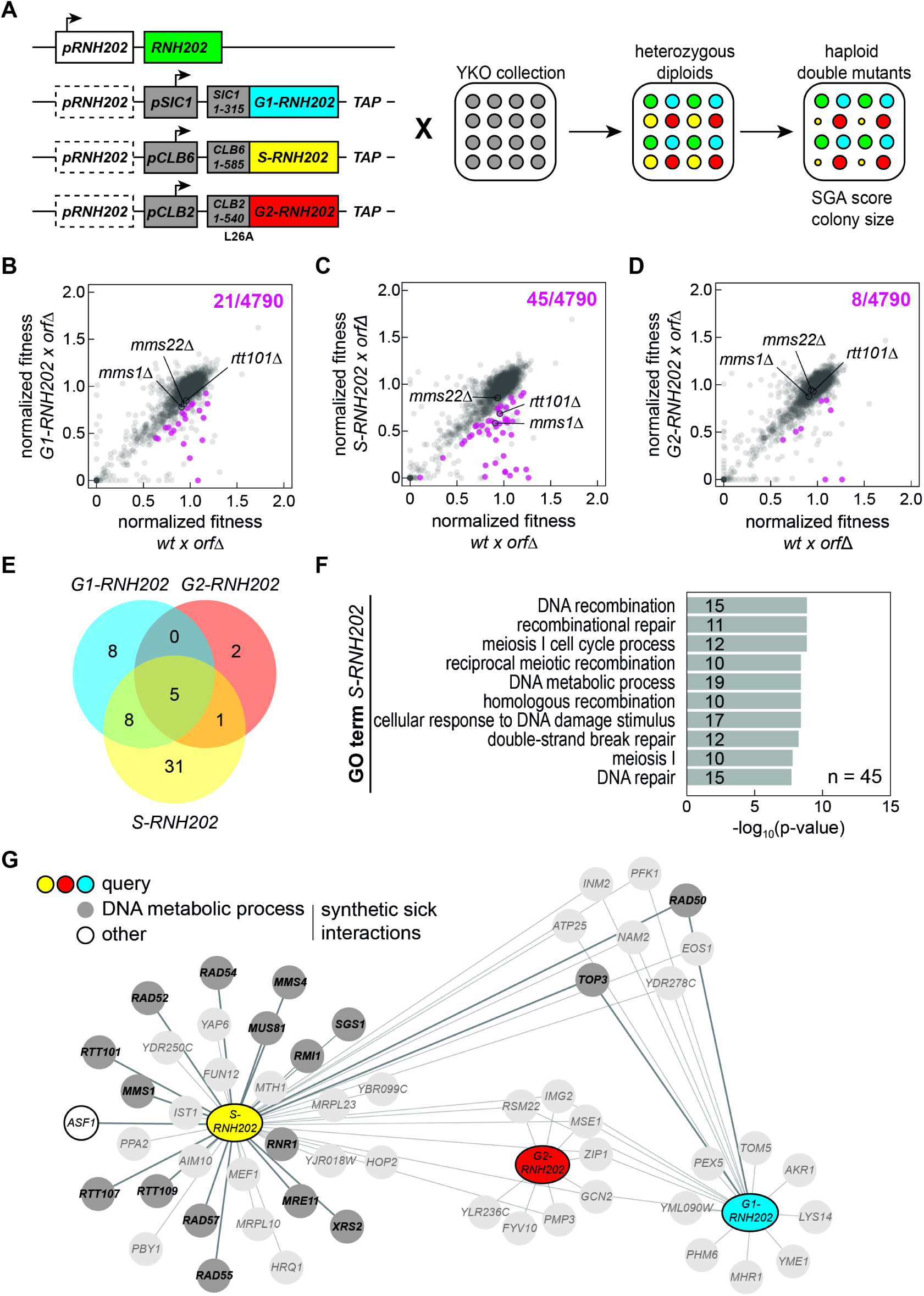
SGA screen identifies network of genes required for rNMP-derived lesion tolerance in S phase. **(A)** For the SGA analysis, the illustrated three query strains (*G1-RNH202*, *S-RNH202*, *and G2-RNH202*) were crossed to the non-essential yeast knockout (YKO) collection. The heterozygous diploids were sporulated and fitness of the resulting haploid double mutants was scored based on their colony size. The outcome was compared to the corresponding scores from the wild type (*RNH202*) cross. For each genotype four replicates per strain were generated and analysed. **(B-D)** Scatter plots of normalized double mutant fitness for the queries compared to wild type (wt). The three queries with cell cycle *RNH202* alleles compared to the wild type control (wt). Each data point represents a single mutant in the YKO collection. Significant synthetic sick interactions (fitness query x *orf*Δ / wt x *orf*Δ < 0.8, p < 0.05 in a t-test, corrected for multiple testing using the Benjamini-Hochberg method) are highlighted in magenta. Crosses with *mms1*Δ, *mms22*Δ and *rtt101*Δ mutants are indicated. Top right, total number of significant synthetic sick interactions. **(E)** Venn diagram of the number of synthetic sick interactions for the three queries with the cell cycle restricted *RNH202* alleles. **(F)** GO term enrichment analysis for synthetic sick interactions of the *S-RNH202* query. Only Biological Process GO terms are shown, top 10 terms by p-value in a hypergeometric test. **(G)** Network summary of the synthetic sick interactions for the three queries with the cell cycle *RNH202* alleles. Genes mapped to the GO term „DNA metabolic process” are highlighted in grey. We could exclude genes with linkage to the *rnh202*Δ locus and manual tetrad dissection identified false positives (***Table S1***). These false positive hits were excluded from the network (faint appearance in the scheme).

In summary, the SGA screen identified 45 candidate genes linked to DNA metabolic processes, including DNA resection, HDR, and repair intermediate resolution, that may be involved in repair of rNMP-derived gDNA lesions in S phase with 31 being unique to this process and 17 confirmed, including HDR.

### Rtt101 acts in a genetic pathway with Rad51 to promote rNMP repair

Genetic evidence points to a crucial role of the Rtt101^Mms1-Mms22^ ubiquitin ligase complex in the regulation of DNA repair and chromatin establishment (Buser et al., 2016; Han et al., 2013; Luke et al., 2006; Mimura et al., 2010; Zaidi et al., 2008). These studies addressed the role of Rtt101 in the presence of exogenously induced DNA damage such as the Top1 poison CPT, the alkylating agent MMS, or the ribonucleotide reductase inhibitor HU. Here, we found that Rtt101and the adaptor subunit Mms1 are also required when endogenous DNA lesions at genomic rNMP arise in S phase (***Figure 1G***). *MMS22* was not a hit in the screen but was manually confirmed (***Figure 2A***).

**Figure 2.**
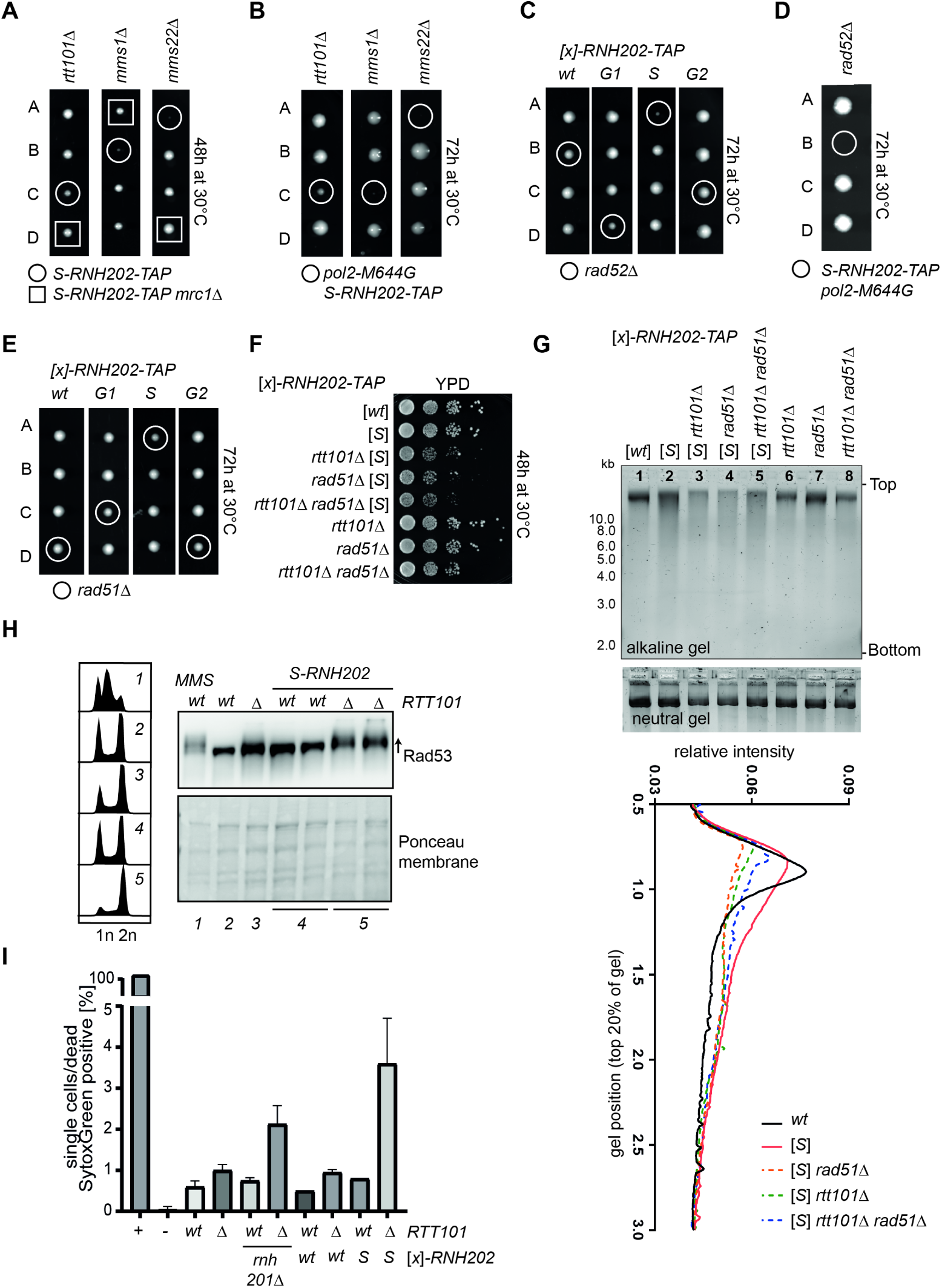
Rtt101, histone modifiers and HDR factors are required to promote cell viability when genomic rNMPs are hydrolysed in S phase. **(A-E)** Diploids were micromanipulated onto rich medium, and the agar plates were grown for the indicated time at 30°C. **(A)** Representative tetrads from the dissection of Rtt101^Mms1-Mms22^ complex-deficient diploid strains in combination with the *S-RNH202-TAP* allele revealed smaller colony sizes after 48h outgrowth at 30°C (colonies in circles). The genetic suppressor of *rtt101*Δ, *mrc1*Δ is sufficient to fully reverse the growth phenotype as shown before for *rtt101*Δ sensitivity in the presence of MMS or CPT (Buser et al., 2016) (colonies in squares). **(B)** Representative tetrads from dissections of Rtt101^Mms1-Mms22^ complex-deficient diploid strains in the *S-RNH202-TAP pol2-M644G* genetic background augmented the sickness. The *rtt101*Δ *S-RNH202-TAP pol2-M644G* lethality was less penetrant compared to the *mms1*Δ and *mms22*Δ mutants, but after propagating, these small colonies were mostly inviable or acquired suppressor mutations. **(C)** Representative tetrad dissections of *rad52*Δ with all cell cycle alleles of *RNH202-TAP* demonstrates that the *S-RNH202-TAP rad52*Δ double mutant is inviable. **(D)** The phenotype from C) was exacerbated when increasing the rNMP load using an *S-RNH202-TAP pol2-M644G* double mutants. This confirms that *S-RNH202-TAP* requires *RAD52* for survival in the presence of high rNMP load in the gDNA. **(E)** The representative tetrads for the deletion of *RAD51* with all cell cycle alleles of *RNH202* shows that the *S-RNH202 rad51*Δ double mutant is sick, but viable, hence it is possible to do experiments analyzing the HDR (Rad52/Rad51) contribution in the *S-RNH202-TAP* background. **(F)** Tenfold serial dilution of the indicated strains was spotted onto YPD agar plates. Images were taken after 2 days of growth at 30°C. The *S-RNH202-TAP rtt101*Δ and *S-RNH202-TAP rad51*Δ double mutants showed the same degree of synthetic sickness on YPD agar plates. The *S-RNH202-TAP rtt101*Δ *rad51*Δ triple mutant strains shows the same degree of sickness as the double mutants indicative of epistasis between *RTT101* and *RAD51* in the presence of rNMP-derived nicks. **(G)** Alkaline gel electrophoresis of the same strains used in F) showed epistasis of the genomic DNA fragmentation between *S-RNH202-TAP rtt101*Δ and *S-RNH202-TAP rad51*Δ double mutant and the *S-RNH202-TAP rtt101*Δ *rad51*Δ triple mutant strains (compare lanes 3, 4, 5). The neutral gel was a control for the purity and integrity of the genomic DNA. The quantification of the DNA smear shows that in contrast to *S-RNH202-TAP* alone, the lack of *RTT101* and/or *RAD51* leads to more fragmentation hence higher genomic rNMP load confirming that Rtt101 and HDR contribute to rNMP repair. **(H)** Western blot analysis of checkpoint status by phospho-shift analysis of Rad53. Membrane staining with Ponceau Red showed equal loading. *RTT101* deletion elicits the DNA damage checkpoint accompanied by cell death when nicks accumulate in S phase in the *S-RNH202-TAP rtt101*Δ double mutant. The MMS treated wild type strain produced a strong Rad53 phospho-shift as positive control indicative of the activated checkpoint. The flow cytometry DNA profiles were measured using Sytox Green labeled cells and showed a strong 2n peak accumulation in line with the activated Rad53 checkpoint. **(I)** Viability analysis of exponential cultures stained with Sytox Green. 4% of the *S-RNH202-TAP rtt101*Δ cells underwent cell death in unchallenged growth in agreement with a checkpoint recovery defect and/or persistent DNA damage. “Plus” is the boiled positive control representing 100% dead hence Sytox green positive cells. “Minus” indicates the unstained wild type control.

Consistent with the entire Rtt101^Mms1-Mms22^ complex being important for rNMP tolerance, the individual deletions of *RTT101*, *MMS1*, and *MMS22* are all compromised for growth in combination with *S-RNH202* (***Figure 2A***). While colony growth is mildly affected in *rtt101*Δ *S-RNH202*, the deletion of *MMS1* and *MMS22* results in stronger effects (***Figure 2A***). Increasing the rNMP load 10-fold using the *pol2-M644G* allele augments the synthetic sickness of *S-RNH202* expression in Rtt101 complex mutants (***Figure 2B***). Although small spores form initially, they eventually become inviable indicating the presence of severe genomic instability in these strains. The fork protection protein and checkpoint regulator Mrc1 is the major suppressor of *rtt101*Δ (Buser et al., 2016). Noteworthy, *mrc1*Δ was sufficient to rescue the growth of Rtt101^Mms1-Mms22^ deficient *S-RNH202* strains (***Figure 2A***). We are following up the underlying mechanism as to why *mrc1*Δ rescues this phenotype, which will be described elsewhere. We hypothesize that loss of Mrc1 can lead to uncoupling of the replisome, releasing single-stranded DNA that can facilitate HDR in the absence of Rtt101-specific pathways such as the one presented here.

Next, we wanted to assess the role of Top1 and R-loops in the synthetic sick interaction of *S-RNH202* and *RTT101*. However, the *rtt101*Δ *S-RNH202 pol2-M644G* mutant is inviable (***Figure 2B***), we switched to HU instead of *pol2-M644G* to modulate the genomic rNMP load. As expected, in the presence of HU, the *S-RNH202* allele was synthetic sick with *rtt101*Δ (***Figure S2A***). When restricting the expression of RNase H2 to S phase we, at the same time, remove canonical RER, which happens mainly in G2 phase (Lockhart et al., 2019). The Top1-mediated RER-backup pathway can be mutagenic and accounts for cellular toxicity in absence of RNase H2 (Kim & Jinks-Robertson, 2017). Strikingly, the synthetic sickness of *rtt101*Δ *S-RNH202* was Top1-independent (***Figure S2B***). We also confirmed by RNase H1 overexpression in the same strains that the Rtt101-dependent role in *S-RNH202* is R-loop independent (***Figure S2C***). Finally, we combined the *S-RNH202* allele with the RER-deficient *rnh201-RED* allele, a separation of function mutant of RNase H2 ((Chon et al., 2013), reviewed in (Cerritelli & Crouch, 2019)), that is RER defective, but can still remove R-loops, to show that hydrolysed rNMPs in S phase require Rtt101 (***Figure S2D***). Together, we showed that the role of Rtt101 in cells with high rNMP load and RNase H2 dysfunction is independent of Top1 and R-loops.

Next, we wanted to test if Rtt101 and HDR genes repair genomic rNMPs and their lesions. To this end we performed genetic epistasis experiments that place the genes into the same pathway and alkaline gel electrophoresis to monitor the genomic rNMP abundance in the presence and absence of the putative repair pathway. As previously demonstrated, the *RAD52* gene becomes essential when rNMPs hydrolyse enzymatically through *S-RNH202* (***Figure 2C***). This phenotype is exacerbated when rNMPs loads are increased through *pol2-M466G* expression (***Figure 2D***). Expression of *G1-RNH202* also slightly affects the growth of *rad52*Δ cells, which again suggests that RER may be occurring in G1, but some unrepaired nicks are carried into S phase (***Figure 2C***). Due to the lethality of *RAD52* deletions in the *S-RNH202* genetic background, we could not perform genetic epistasis experiments with loss of *RTT101*. The *rad51*Δ *S-RNH202* double mutant, however, is growth impaired to the same degree as the *rtt101*Δ *S-RNH202* double mutant (***Figure 2E***). The *rtt101*Δ *rad51*Δ *S-RNH202* triple mutant is not additive, suggesting that *RTT101* and *RAD51* may function in the same genetic pathway of rNMP- derived nick repair in S phase (***Figure 2F***). We employed alkaline gel electrophoresis to visualize the genomic rNMP load in *rtt101*Δ and *rad51*Δ strains in the presence of increased S phase rNMP-nicking (*S-RNH202*) (***Figure 2G***). In addition to rNMP-hydrolysis activity in S phase, the *S-RNH202* strain lacks canonical G2 phase RER. Therefore, higher rNMP load in the *S-RNH202* strain compared to the wild type *RNH202* allele was expected (***Figure 2G***, lane 1 compared to lane 2). Strikingly, the loss of either *RTT101* or *RAD51* alone, and in combination, resulted in higher DNA fragmentation in alkaline conditions and loss of the prominent genomic DNA band, indicative of fragmented genomic DNA, hence lack of rNMP repair (***Figure 2G***, lanes 3, 4, 5 compared to 2, and quantification graph). Rtt101 deficiency is characterized by a slower checkpoint recovery (Luke et al., 2006). Hence, *rtt101*Δ strains show a broadened 2n peak in DNA profiles (***Figure 2H***, DNA profile 3) and basal checkpoint activation visualized by phospho-Rad53 analysis (***Figure 2H***). In line with the elevated rNMP load and non-repaired DNA damage leading to impaired viability, the *rtt101*Δ *S-RNH202* mutants have fully activated the Rad53-checkpoint (***Figure 2H***). In addition, we measured 4% cell death in that population without further challenge (***Figure 2I***). This supports the idea of a repair pathway as loss of the repair factors, Rtt101 and Rad51, result in a repair defect accompanied by an activated DNA damage checkpoint, hence the failure to efficiently remove rNMPs. Together, these data demonstrate that the Rtt101 complex works together with the recombination machinery to repair rNMPs, and not R-loops, that get nicked in the S phase.

### Rtt101 becomes essential in S phase to overcome Top1-independent rNMP-derived toxicity

We have demonstrated that the *S-RNH202* allele is very similar to the RNase H2 deletion (***Figure S1A-S1C***), hence we predicted that *RTT101* would also play an important role in the S phase repair of hydrolysed rNMPs in RER-deficient strains. Indeed, the *rtt101*Δ *rnh201*Δ double mutants were highly sensitive to HU as compared to the respective single mutants (***Figure 3A)***. Importantly, *RNH1* overexpression, which reduces R-loop levels, did not rescue the *rtt101*Δ *rnh201*Δ viability defect in the presence of HU (***Figure 3A***), suggesting that R-loops may not be responsible for the growth defects. As RNase H2 has a dual role in RNA-DNA hybrid removal, and participates in R-loop removal (Cerritelli & Crouch, 2009; El Hage et al., 2014), we again employed the *rnh201-RED* allele that retains R-loop removal activity but fully lacks RER-activity ((Chon et al., 2013), reviewed in (Cerritelli & Crouch, 2019)). We found that the RER-proficient *RNH201* wild type allele could rescue the growth defect of *rtt101*Δ *rnh201*Δ mutants in the presence of HU, however strains expressing the *rnh201-RED* allele were as sick as the vector control (***Figure 3B***). This confirmed that persisting genomic rNMPs are the underlying cause of the slow growth in *rtt101*Δ *rnh201*Δ cells (***Figure 3A, 3B***). The *rtt101*Δ *rnh201*Δ *pol2-M644G* triple mutant was genetically unstable, therefore we employed an *RNH201-AID\** auxin-inducible degron (Morawska & Ulrich, 2013), to highly reduce RNase H2 activity in the presence of auxin (***Figure S3A, S3B***). Similar to the *rtt101*Δ *rnh201*Δ double mutant, *rtt101*Δ *RNH201-AID\** cells presented a mild growth defect upon exposure to auxin (***Figure 3C***). Upon addition of the *pol2-M644G* allele to increase the genomic rNMP load, the *rtt101*Δ *RNH201-AID* pol2-M644G* triple mutant was inviable in the presence of auxin (***Figure 3C***). The *rnh201-RED* allele could not rescue the synthetic lethality of the *rtt101*Δ *RNH201-AID* pol2-M644G* triple mutants in the presence of auxin (***Figure 3D***). The deletion of both *MMS1* and *MMS22* showed similar genetic interactions with *RNase H2* impairment, suggestive of the entire E3 ubiquitin ligase complex being required to tolerate increased rNMP levels (***Figure S3C***). As with the *S-RNH202 allele*, we asked whether Top1 mutagenesis was responsible for the severe phenotype of *rtt101*Δ *RNH201-AID* pol2-M644G* cells. In line with the *S-RNH202 allele* (***Figure S2B***), the deletion of *TOP1* did not rescue the viability of *rtt101*Δ *RNH201-AID* pol2-M644G* in the presence of auxin (***Figure 3E***). This was consistent for the entire Rtt101^Mms1-Mms22^ complex (***Figure S3D***). As we previously demonstrated that *RTT101* acts in the same pathway as HDR for survival with *S-RNH202* expression, we also expected that defective RER would lead to a fitness disadvantage when HDR was inactive. To this end, we observed that the loss of *RAD52* was defective for growth in the presence of the *rnh201-RED* allele (***Figure S3E***). The viability of a RER-deficient *pol2-M644G* strain fully relied on the presence of *RAD52*, furthermore indicating that the lesion potential correlates directly with the amount of rNMPs (***Figure S3F***). Together, these results are consistent with an Rtt101-mediated HDR being required to repair nicked rNMPs in S phase.

**Figure 3.**
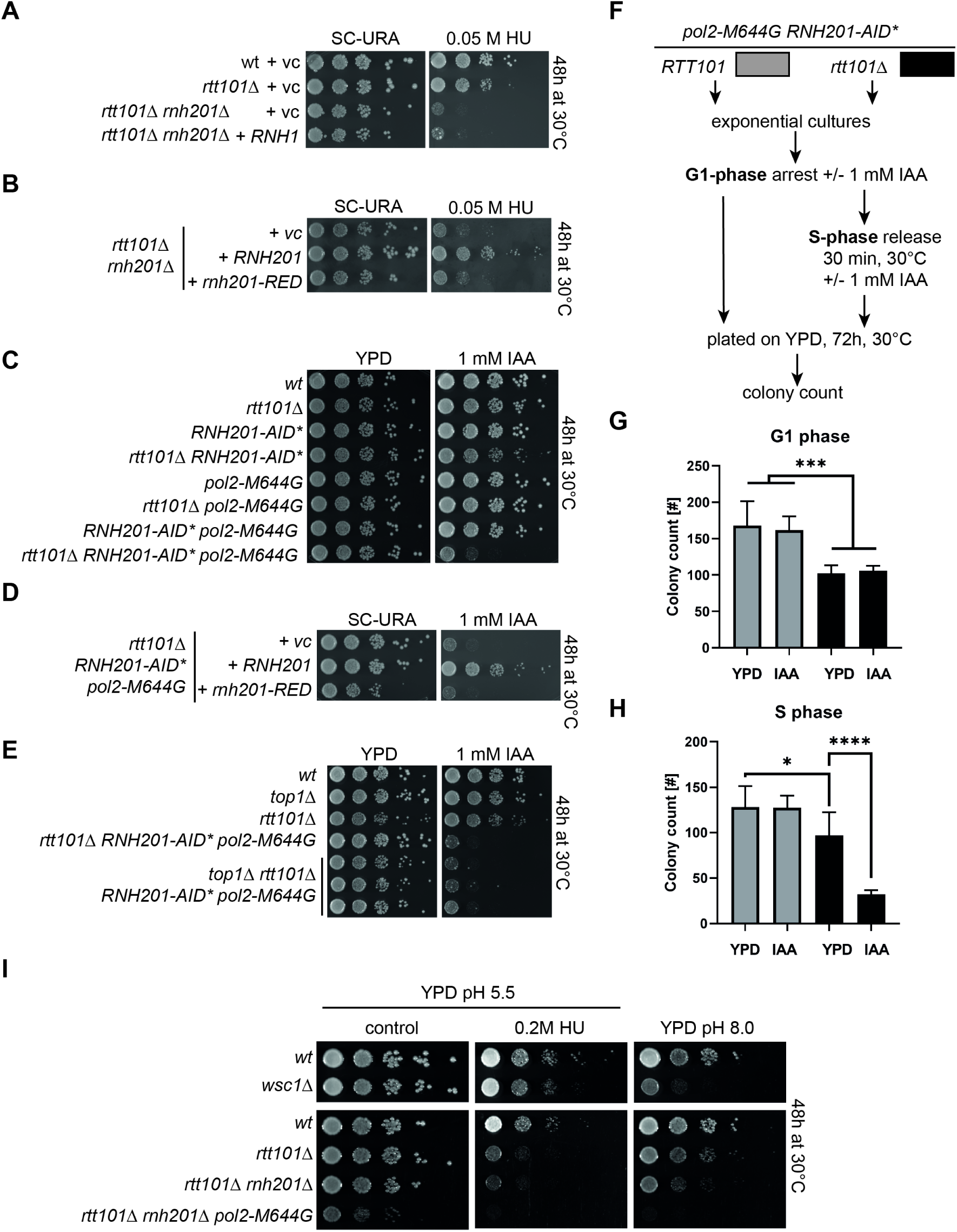
Rtt101 becomes essential in S phase in a Top1-independent manner to overcome rNMP-derived toxicity. **(A-E)** A tenfold serial dilution of the indicated strains was spotted onto the indicated agar plates. Images were taken after 2 days of growth at 30°C. vc = vector control, *RNH1*=RNase H1, hydroxyurea (HU), *RED* allele= ribonucleotide excision deficient (P45D Y219A) (Chon et al., 2013), IAA = indole acetic acid (auxin). **(A)** The *rtt101*Δ *rnh201*Δ double mutant is synthetic lethal in the presence of HU. Transformation with *RNH1* did not affect the growth of the double mutant on HU containing agar plates. **(B)** Transformation with wild type *RNH201* did rescue growth of the *rtt101*Δ *rnh201*Δ double mutant on HU plates, while the RER-deficient separation-of-function *rnh201-RED* mutant had no effect. **(C)** Depletion of *RNH201-AID\** in the presence of auxin resulted in a synthetic sick growth phenotype with *rtt101*Δ, which was amplified into a synthetic lethal phenotype when combined with *pol2-M644G*. **(D)** Complementation of the *pol2-M644G rtt101*Δ *RNH201-AID*\* triple mutant with wild type *RNH201* rescued growth on auxin plates, while the *rnh201-RED* mutant did have no effect. **(E)** The synthetic lethality of the *pol2-M644G rtt101*Δ *RNH201-AID\** triple mutant on auxin plates was Top1-independent. Three independent strains from separate tetrads were spotted to confirm Top1-independence of the observed phenotype. **(F-H)** Liquid cultures with the indicated genotypes were synchronized with α-factor in the G1 phase in the presence of auxin to deplete *RNH201-AID\**. The arrested cultures were either directly plated on YPD agar plates (colony count shown in panel (G)), or released into the S phase in the presence of auxin, followed by plating on YPD agar plates (colony count shown in panel (H)). The colony formation capacity was assessed by counting the colonies after 72h growth at 30°C. Bar graph are summarizing n=7 plate counts per genotype and condition. Data is represented with mean with SD. Statistics were performed with GraphPad Prism8. Unpaired, two-tailed student t test; ****p<0.0001; ***p<0.001; *p<0.05. **(I)** The serial dilution assay with *rtt101*Δ strains in the RER-deficient *rnh201*Δ and the rNMP accumulating *pol2-M644G* background revealed that alkalization of the YPD agar was sufficient to phenocopy the synthetic sick growth defects seen in the presence of HU. Images were taken after 2 days of growth at 30°C.

The Rtt101^Mms1-Mms22^ ubiquitin ligase complex is associated with the replisome during S phase (Buser et al., 2016) and becomes essential when rNMPs are hydrolysed in S phase by *S-RNH202* (***Figure 1, 2***). We wanted to test if rNMP-derived damage in a single S phase requires the immediate activity of Rtt101^Mms1-Mms22^. Therefore, we performed a colony formation assay to assess cell viability when rNMP removal is prevented either in the G1 phase or in G1 phase and during S phase entry and progression (***Figure 3F-H, S3G-H***). We arrested *RNH201-AID* pol2-M644G* and *rtt101*Δ *RNH201-AID* pol2-M644G* cultures in the G1 phase in the presence of auxin to degrade Rnh201 and prevent RNase H2 activity. To assess the toxicity of rNMP accumulation in G1 phase, we plated the cultures directly on rich medium and quantified the number of colonies formed (***Figure 3G***). Alternatively, the synchronized cultures were released from the G1 arrest into the S phase still in the presence of auxin to abolish RNase H2 activity during S phase entry and progression. S phase cultures were also plated on rich medium, thereby allowing the re-accumulation of RNase H2 (***Figure 3H***). We monitored the cell cycle phases of the cultures by flow cytometry (***Figure S3F***). We observed an overall 20% viability reduction in the *rtt101*Δ background (***Figure 3G, 3H***). RER-deficiency did not affect the cell viability during G1 phase (***Figure 3G***, compare black columns). However, in the absence of Rtt101 there was a 70% reduction in cell viability when RER deficient cells progressed through S phase (***Figure 3H***, compare black columns). Therefore, non-repaired rNMPs are only toxic in *rtt101*Δ cells in the S phase of the cell cycle, and not in G1.

The 2’-hydroxyl group renders rNMPs susceptible to spontaneously hydrolyse the phosphodiester backbone compared to the more stable and resistant DNA deoxy sugars. Since this hydrolysis reaction is more likely in a basic environment, we assumed that growth in alkaline conditions may increase the likelihood that hydrolysis at genomic rNMPs will occur. Alkaline conditions were therefore expected to impact the growth of *rtt101*Δ strains similar as the presence of hydroxyurea or the absence of RER. *Saccharomyces cerevisiae* media (YPD) has pH5.5 and therefore is mildly acidic. We increased the pH of solid agar medium to pH8.0. We confirmed alkaline pH8.0 in agar plates by using the *wsc1*Δ strain that renders cells sensitive to alkali pH stress (Serra-Cardona et al., 2015) (***Figure 3I***). The *rtt101*Δ strain was mildly sensitive to pH8.0 whereas *rtt101*Δ *rnh201*Δ cells were highly sensitive to alkaline conditions (***Figure 2I***). The unstable *rtt101*Δ *rnh201*Δ *pol2-M644G* triple mutant was fully inviable on pH8.0 (***Figure 2I***).

In summary, we report that the negative genetic interaction between the deletion of Rtt101^Mms1-Mms22^ ubiquitin ligase subunits and RNase H2 defects is due to RER-deficiency and is exacerbated in rNMP accumulating (*pol2-M644G*, HU) condition. In RER-defective cells, rNMPs are likely hydrolysed prior to, or during, S phase and require Rtt101 mediated HDR for repair upon encounter with the replisome. In line with the physical association with the replisome in S phase (Buser et al., 2016), Rtt101 function is essential during S phase to counteract rNMP-derived cellular toxicity.

### Rtt101 mediates the repair of rNMP-derived DNA damage in S phase through histone H3 ubiquitylation

Using different genetic models (*S-RNH202* allele, RNase H2 deletion, alkaline conditions, *pol2-M644G* allele, *rnh201-RED* allele), we demonstrated that the Rtt101^Mms1-Mms22^ ubiquitin ligase complex is required to deal with Top1-independent rNMP-derived DNA damage in S phase. We speculate that we may have found the genetic requirements for a unique rNMP-derived lesion repair pathway that acts in S phase, complementing the G2 phase RER and the Top1 pathways (Kellner & Luke, 2020). We set out to get a deeper molecular understanding by further probing the genetic interactions from the *S-RNH202* SGA genetic network (***Figure 1***) and potentially identify substrates for Rtt101.

In general, the identification of ubiquitin ligase substrates has proven to be challenging because the cullin enzymes are scaffolds forming various multi-protein complexes (Finley et al., 2012). Additional hurdles include the characterisation of ubiquitin-modified substrates due to the plethora of consequences ubiquitylation inflicts, i.e. proteasomal degradation, signaling, conformational change, protein-protein interaction changes (García-Rodríguez et al., 2016). The Rtt101^Mms1-Mms22^ complex has previously been shown to ubiquitylate histone H3 on three lysine (K) residues (K121, K122 and K125) (Han et al., 2013). The modification does not lead to proteasomal degradation, but rather facilitates the deposition of newly synthesized histones during replication-coupled nucleosome assembly (RCNA). Other, non-replication related, substrates of Rtt101^Mms1-Mms22^ have been reported in yeast (Han et al., 2010), whereas multiple targets of Cul4 have been elucidated in human cells (Higa et al., 2006; Liu et al., 2019; Thirunavukarasou et al., 2014; Ye et al., 2019; Zhao et al., 2010; Q. Zhu et al., 2017).

The histone chaperone Asf1 and the histone acetylase Rtt109, which acetylate lysine 56 of H3 (H3K56), act upstream of Rtt101^Mms1-Mms22^ in terms of nucleosome assembly (Han et al., 2013) (***Figure 4A***). Similar to *RTT101,* the *ASF1* and *RTT109* genes were essential for cellular survival when rNMPs were nicked in the S phase (***Figure 4B***). In the RCNA pathway, Rtt101 is responsible for the ubiquitylation of newly synthesized histone H3 on three lysine residues, which release the H3-H4 dimer from the histone chaperone Asf1 (***Figure 4A***). As the downstream RCNA factors *CAC1/RLF2*, *CAC2*, *CAC3/MSI1* that form the CAF-1 complex and *RTT106* have redundant roles (Clemente-Ruiz et al., 2011), the single deletions do not affect *S-RNH202* colony growth (***Figure 4C***). CAF-1 deletion combined with *RTT106* deletion is synthetic lethal, which is why we cannot rule out their contribution. We generated heterozygous diploid strains and derived the haploid double mutants to test the impact of Rtt109 and Asf1 loss in a RER-deficient condition using the *RNH201-AID\** degron and the *pol2-M644G* allele (***Figure 4D, 4E***). Strikingly, triple mutants displayed lethality in the presence of auxin, reflecting the major role of RCNA factors, Asf1 and Rtt109, in the repair of rNMP-derived lesions (***Figure 4D, 4E***).

**Figure 4.**
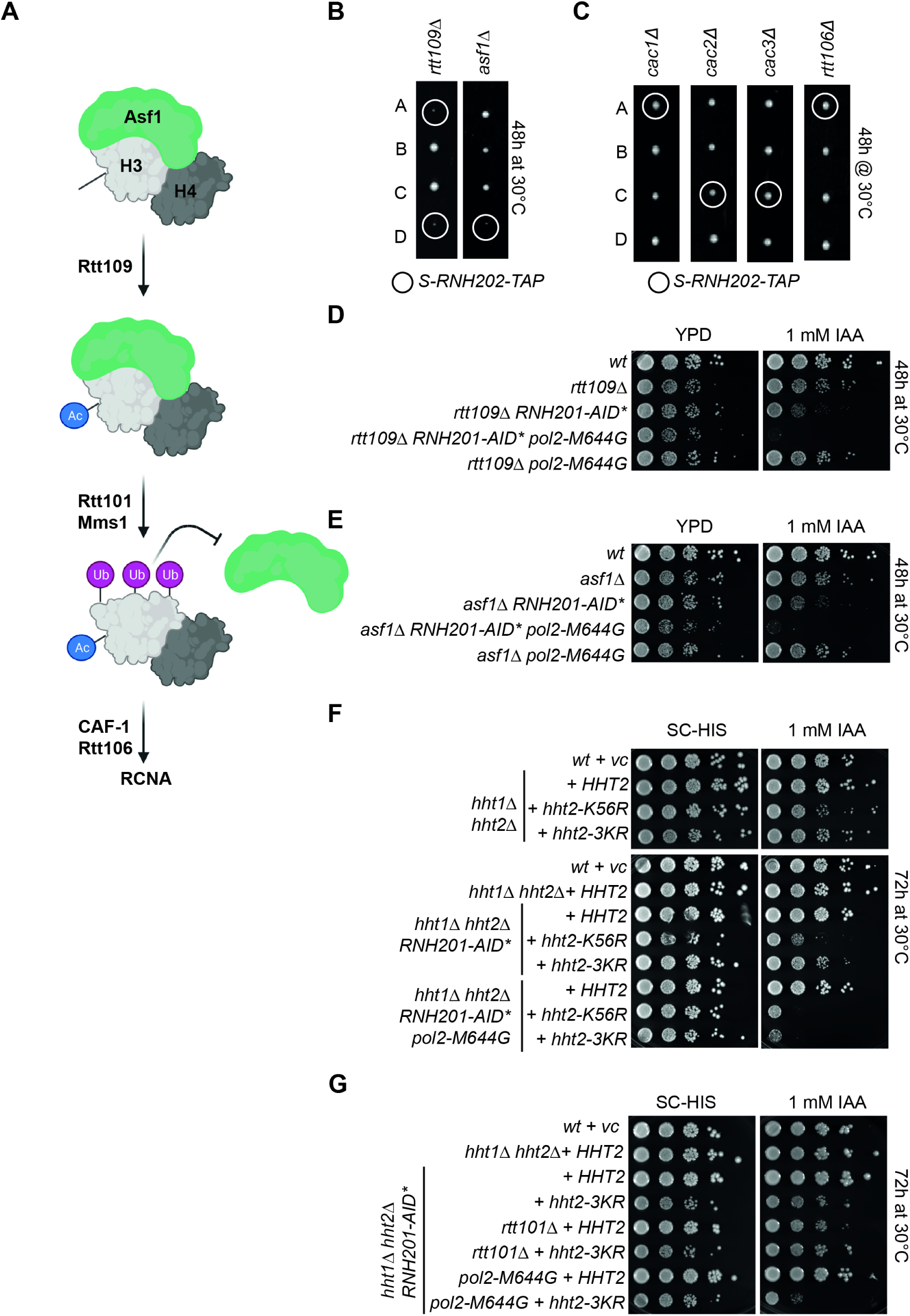
Rtt101 mediates the repair of rNMP-derived DNA damage in S phase through histone H3 ubiquitylation. **(A) S**cheme of the contribution of the histone chaperone Asf1, the histone acetyl transferase Rtt109 and the ubiquitin ligase Rtt101^MMS1^ to the replisome coupled nucleosome assembly (RCNA) pathway; **i)** sequence of events (modified from (Han et al., 2013)): Asf1 binds to de novo synthesized H3-H4 dimers and Rtt109 acetylates H3 followed by ubiquitylation of H3 through Rtt101 that lead to the release of Asf1 and facilitates DNA incorporation; **ii)** acetylation deficient *K56R* mutant is reminiscent of *RTT109* deletion; **iii)** ubiquitylation-deficient *3KR* mutant is reminiscent of the *RTT101* deletion. **(B)** Manual tetrad dissection confirmed the synthetic lethal phenotype between the *S-RNH202* allele and the histone remodeler genes *RTT109* and *ASF1* (double mutant colonies in circles). **(C)** Representative tetrads from single CAF-1 complex deletion mutants (*cac1*Δ, *cac2*Δ, *cac3*Δ) in combination with the *S-RNH202* allele to check contribution of the RCNA pathway. Note that these genes work redundantly and they are synthetic lethal with each other, hence we cannot exclude their contribution. **(D)** Serial dilution assays with the histone acetylase *Rtt109* deletion mutants shows that loss of *RTT109* is toxic in RER-deficient strains with high rNMP load (*pol2-M644G*). **(E)** The same is true for loss of the histone chaperone *ASF1*. **(F)** Serial dilution spot assays with histone 3 mutants deficient for Lysine-56 acetylation (“H3-K56R”) and Rtt101-dependent Lysine-121,122,125 triple ubiquitylation (“H3-3KR”) (Han et al., 2013). These plasmid-borne mutant versions of histone H3 replaced the *HHT1* and *HHT2* that were deleted. Histone 3 Lysine-56 acetylation became essential in RER-deficient cells (*RNH201-AID\** on IAA plates). The H3-3KR strain revealed mild sickness in RER- deficient cells but was inviable when rNMP levels increased with the pol2-M644G allele. IAA = indole acetic acid (auxin) **(G)** Strains from (F) were combined with *RTT101* deletion to confirm the epistasis between *RTT101*-deficiency and H3 ubiquitylation deficiency (compare lanes 4-6).

To assess if the Rtt101-dependent H3 ubiquitylation has a direct role in rNMP-lesion repair, we combined the ubiquitylation-deficient H3-3KR mutant (Han et al., 2013) with an RER-deficient background using the *RNH201-AID\** degron (***Figure 4F***). Rtt109-mediated H3-K56 acetylation occurs upstream of Rtt101-dependent H3 ubiquitylation (Han et al., 2013). The H3-K56R acetylation-deficient strain was synthetic sick with loss of RNase H2 and inviable when rNMPs accumulate in the *RNH201-AID* pol2-M644G* strain background (***Figure 4F***). Interestingly, the H3-3KR ubiquitylation-deficient mutant could support growth upon loss of RNase H2 better than the H3-K56R mutant, however, Rtt101- dependent H3-ubiquitylation became essential when rNMP load increased in the *RNH201-AID* pol2-M644G* strain (***Figure 4F***). To show that Rtt101 and the Rtt101-dependent H3 ubiquitylation behave in an epistatic manner, we deleted *RTT101* in the *H3-3KR* mutant strains. Indeed, deleting *RTT101*, or impairing H3 ubiquitylation (*H3-3KR*), or the combination of both impaired cell viability to the same degree in RER-deficient strains (***Figure 4G***). This suggests that histone H3 is a key target of Rtt101 and we conclude that Rtt101^Mms1-Mms22^ dependent histone H3 ubiquitylation at lysines-121, -122, and -125 is critical for the repair of rNMP-derived DNA damage. However, it also suggests that other functions of the RCNA pathway may be important for rNMP repair in RER defective strains. Interestingly, Asf1 has a role in the regulation of Rad53 checkpoint control (Tsabar et al., 2016). It will be important to unravel the unknown connections that still exist between Rtt101, DNA repair, checkpoint recovery, and nucleosome RCNA.

## Discussion

Eukaryotic cells repair genomic rNMPs by RNase H2-initiated ribonucleotide excision repair (RER). The loss of RNase H2 function leads to the accumulation of genomic rNMPs, which then become to an extent substrates for error-prone repair by topoisomerase 1 (reviewed in (Kellner & Luke, 2020)). Recently, it has been demonstrated that some human cancers harbor RNase H2 mutations, resulting in rNMP accumulation and Top1-mediated genome instability (Zimmermann et al., 2018). These cancers are considered “druggable” as Top1 lesions recruit PARP to sites of damage and hence become susceptible to PARP inhibitors (Zimmermann et al., 2018). Elucidating alternative rNMP repair pathways may yield additional factors and pathways that could potentially be targeted in RER defective human cancer cells. Importantly, it has been shown that RER-defective budding yeast have a nearly identical mutagenic signature profile as RER defective cancer cells (Reijns et al., 2022), hence making yeast a highly relevant model for the study of rNMP repair.

The loss of *RAD52* becomes essential in RER-defective yeast cells and *TOP1* deletion can only partially rescue the loss of fitness, indicating that there might be additional sources for rNMP-mediated genome instability apart from Top1 (Huang et al., 2017; Lockhart et al., 2019). Genomic rNMPs are prone to hydrolysis and nick formation and it was shown recently that the CMG helicase will eventually run off the DNA, if the leading strand template is nicked upstream of the replication fork (Vrtis et al., 2021). Hence, we hypothesized that the HDR machinery was repairing rNMP-induced lesions (seDSB) that occur when a nicked rNMP encounters replication (Lockhart et al., 2019). In support of this idea, increased rNMP-nicking in RER-deficient cells rendered cells dependent on HDR, independent of Top1 (Lockhart et al., 2019).

Here, we employed the *S-RNH202* allele to induce seDSBs at rNMPs to look for mutants with reduced fitness similar to *rad52*Δ, in a genome-wide screening approach (***Figure 1***). As a result, we elucidated a genetic network for rNMP-derived <u>n</u>ick lesion repair (NLR) (***Figure 5***). NLR includes the Rtt101 ubiquitin ligase, the Rad52-based HDR machinery, the MRX (Mre11-Rad50-Xrs2) complex, and the Rtt109/Asf1 replication-coupled nucleosome assembly (RCNA) pathway (***Figure 1***). In addition, we found the RST (Rmi1-Sgs1-Top3) and Mus81-Mms4 complexes, which likely provide resolution of the multiple recombination intermediates formed during HDR (Hickson & Mankouri, 2011) (***Figure 1***). We showed that the exclusive nicking of rNMPs in S phase is particularly toxic in *rtt101*Δ cells (***Figure 2A***). Indeed, we could demonstrate that loss of *RAD51* and *RTT101* are epistatic in terms of rNMP repair (***Figure 2F, G***). Furthermore, we confirmed that loss of *RTT101* was sufficient to kill RER-deficient cells in a Top1-independent manner (***Figure 3E***). We were also able to conclude that Rtt101 function is required in S phase (***Figure 3H***), which is in alignment with its replisome association (Buser et al., 2016). Moreover, the mutated allele of histone H3 that can no longer be ubiquitylated by Rtt101 (H3-3KR) also renders cells highly sensitive to high levels of rNMPs (***Figure 4F-G***). Although Rtt101, HDR and H3 ubiquitylation are all working together in a genetic pathway it remains unclear as to how Rtt101 promotes HDR.

**Figure 5.**
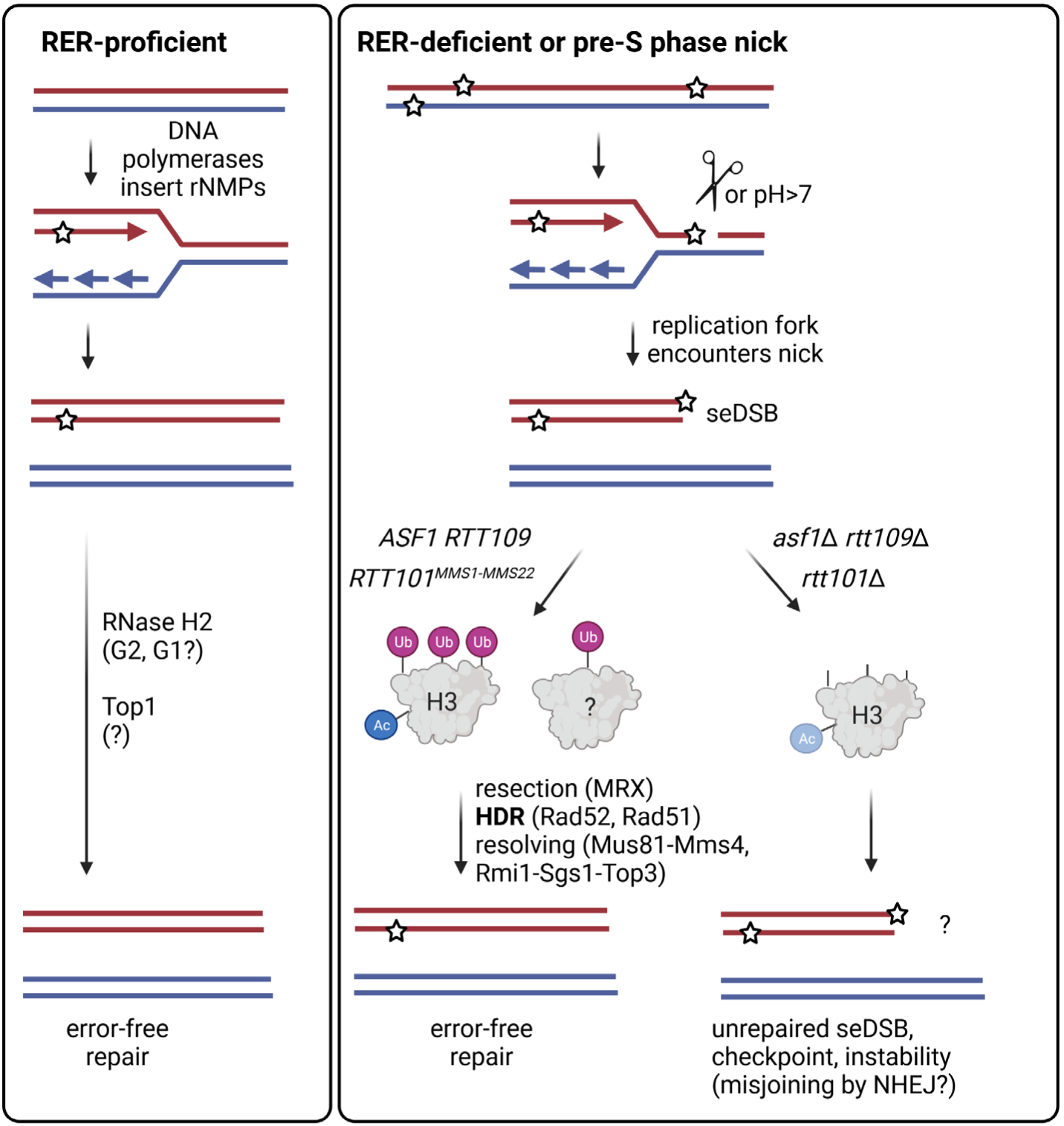
Model Top1-independent NLR pathway is essential when rNMPs cause pre-S phase nicks that result in seDSB. DNA polymerases transiently incorporate single rNMPs into the genome during replication and repair. The RER pathway removes genomic rNMPs immediately in the subsequent G2 phase. In RER-deficient, or RER-dysfunctional cells the Top1-mediated backup pathway deals with rNMP-removal. However, if high amounts of genomic rNMPs accumulate in RER-deficient, or RER-dysfunctional cells, the likelihood increases that hydrolysis-prone rNMPs form ssDNA nicks. When the replication fork encounters such an rNMP-derived nick in the leading strand template, a toxic seDSB is formed. To repair the rNMP-derived seDSB lesions, functional RCNA is required. The histone remodelers Asf1 and Rtt109 act upstream of Rtt101^Mms1-Mms22^, presumably accompanied by the resection of the seDSB by MRX (Mre11-Rad50-Xrs2), followed by HDR (Rad52, Rad51) and resolution of the HDR intermediates (Mus81-Mms4, Rmi1-Sgs1-Top3) to result in the error-free repair of the seDSB. In *RTT101*-deficient cells with high rNMP load, histone H3 remains does not become ubiquitylated and downstream error-free HDR repair of the seDSB is compromised causing genomic instability likely by alternative, error-prone repair attempts. Abbreviations: NLR = rNMP-derived nick lesion repair, rNMP = single ribonuclesoide monophates, RER = ribonucleotide excision repair, seDSB = single-ended double strand break, ssDNA = single stranded DNA, RCNA = replication-coupled nucleosome assembly, HDR = homology -directed repair

One possibility would be that rNMPs are more susceptible to induce nicks because the chromatin structure of *rtt101*Δ cells is altered due to the RCNA defects. This would be consistent with decreased nucleosome deposition and a more open chromatin state. In agreement, it has been reported that telomeric heterochromatin is lost in *rtt101*Δ and *mms1*Δ mutants (Mimura et al., 2010). It will be important determine if hydrolysed rNMPs are more frequent in more accessible chromatin environments and if such environments actually increase in the absence of the Rtt101 complex. Alternatively, it could be that Rtt101-mediated H3 modification are important for the HDR reaction itself. This hypothesis is supported by the fact that deletion of the fork protection protein and damage checkpoint mediator *MRC1* can rescue the sensitivity of *rtt101*Δ cells to genotoxic agents (Buser et al., 2016) as well as to accumulation of rNMPs (***Figure 2A***). Indeed, Mrc1 can differentially regulate resection and HDR at DSBs (Alabert et al., 2009) and it was recently demonstrated that this involves changes in chromatin compaction (Xing et al., 2021). Further support that the repair of rNMP-derived lesions is coupled to alterations of chromatin was shown in a recent study in human cells (Nakamura et al., 2021). Specifically, they looked at seDSB damage caused by the replisome running into TOP1-DNA adducts after CPT treatment. In line with our yeast genetic network, HDR factors, MRN, RAD51, and MMS22L-TONSL were found to be associated with the broken forks in human cells (Nakamura et al., 2021). In addition, broken and stalled replication forks presented a distinct chromatin environment with a defect in histone deposition (Nakamura et al., 2021).

In addition, sister chromatin cohesion is important at seDSBs to ensure that repair occurs primarily from the sister chromatid and not a homologous chromosome. Rtt101, Mms1 and Mms22 promte sister chromatid cohesion through their replisome association (Zhang et al., 2017). Interestingly, the cohesion- like Smc5/6 complex becomes essential in the absence of RER ((Lafuente-Barquero et al., 2017)) and may also be intertwined with the Rtt101, Rtt109, HDR-mediated repair of rNMPs. In fission yeast, the mega-nuclease complex MRN (Mre11-Rad50-Nbs1) is critical to control sister chromatid cohesion at replication-associated seDSBs to allow HDR repair and prevent Ku-mediated DSB repair (M. Zhu et al., 2018). In accordance, all subunits of the *Saccharomyces cerevisiae* MRX complex seem to be essential for NLR (***Figure S1I***). Since cohesin and cohesin-like factor are essential, they were not revealed in the above-described screen and will have to be tested using temperature-sensitive mutant versions.

It will be important to determine whether the Rtt101 E3 human equivalent, Cullin-Ring-Ligase 4 (CRL4), also contributes to rNMP repair in RNase H2 defective cells, as this may represent alternative therapeutic opportunities, in addition to PARP inhibitors in RER defective cancer cells. It is feasible to put this to the test in the future as the CRL neddylation inhibitor MLN4924 was extensively studied and went into clinical trials for cancer intervention (Aubry et al., 2020; Shah et al., 2016). In this respect it is interesting that the cullin subunit of CRL4 (CUL4A) is overexpressed in many human cancers (Sharma & Nag, 2014). The cancer-specific overexpression is a result of the genomic locus in human cells that undergoes amplification in cancers (Chen et al., 1998). Hence, it is possible that this overexpression promotes CRL4-dependent DNA repair also in the context of other human deficiencies (e.g. RER). We speculate that the role for NLR could be greater than expected as the cytoplasm of a cancer cell is slightly alkaline (pH>7), and would therefore promote rNMP-mediated hydrolysis of the DNA backbone. The intracellular alkalization of cancer cells seems connected to the initial oncogenic transformation and the progression of the tumour (Harguindey et al., 2005; Neri & Supuran, 2011). Translational studies will show if RER-defective human cancer cells with alkaline intracellular environment may even favor NLR due to augmented spontaneous rNMP hydrolysis.

## Materials and Methods

### Yeast strains and plasmids

*Saccharomyces cerevisiae* strains used in this study derive of the standard S288C (*MATa his3*Δ*1 leu2*Δ*0 ura3*Δ*0 met15*Δ*0*) strain and are listed in ***Table S2***. Strains were grown under standard conditions in YPD ( 1% [w/v] yeast extract, 2% [w/v] peptone supplemented with 2% glucose) or in SC (0.2% [w/v] Synthetic Complete medium without specific amino acids, 1% [w/v] yeast nitrogen base supplemented with 2% glucose) at 30°C if not indicated otherwise. Yeast transformations with plasmid or PCR products were performed with the standard lithium acetate polyethylene glycol (PEG) method (Gietz & Woods, 2002). Plasmids and oligonucleotides are listed in ***Table S3***.

### Yeast tetrad dissection

For analysis of the meiotic product, we crossed a *MATa* with a *MATalpha* haploid strain, selected for diploids based on auxotrophy or antibiotic resistance, and patched the diploid strain on rich pre-Sporulation plates (YP agar with 6% [w/v] glucose]. Then we froze part of the patch and transferred part of the patch into Sporulation medium (1% potassium acetate, 0.005% zinc acetate buffer) and incubated the cultures with shaking at 23°C. After a few days, the sporulation cultures were treated in a ratio of 1:1 [v/v] with Lyticase (L4020 Sigma Aldrich, 2.5 mg/ml, 200 units/µl, in 1M D-Sorbitol) to digest the ascus. After 15-20 min at room temperature, the culture was applied to an agar plate and tetrads were dissected using a Singer micromanipulator. Colonies of haploid spores grew at 30°C for three days. Images were taken at 48 and 72h with the ChemiDoc™ Touch Imaging System (Bio-Rad). After three days, the spores were replica plated, genotypes were scored and strains were frozen in 15% glycerol containing cryopreserved stocks at -80°C. Strains are listed in ***Table S2***.

### Flow cytometry analysis for DNA content

Cells were fixed in 70% ethanol overnight and then treated with 0.25 mg/ml DNase- and Protease-free RNase A (ThermoFisher Scientific, 10753721) at 37°C for 2h and Proteinase K (Biofroxx, 1151ML010) at 50°C for 2h in 50 mM Tris-HCl pH7.5 buffer. The cell suspension was sonified using a Branson sonifier 450 for 5 sec with output control 1 and duty cycle constant. Then, cells were stained with a final concentration of 2.4 µM SYTOX Green nucleic acid stain (ThermoFisher Scientific, 1076273). Measurement was performed on the BD LSRFortessa flow cytometer (BD Biosciences) using the BD FACSDiva software (v9.0.1). With low flow rate, 20,000 events were recorded. Analysis was performed with FlowJo (v10.8.0) using the following gating strategy: From the main population in FSC-A vs. SSC-A, doublets were excluded in the Sytox-Green A vs. W channel, and DNA content was assessed in the histogram of the Sytox-Green-A channel (Ex 488nm, 530/30BP).

### Flow cytometry analysis for cell viability

Cells were collected and the cell pellet was washed with 50 mM Tris pH 7.5 and resuspended in 1 ml 50 mM Tris pH 7.5 containing 0.5 μM SYTOX Green. Measurement and analysis were the same as for the DNA content analysis except for doublet exclusion, which was done in the SSC-A vs. W channel. As a control sample for dead cells, controls were incubated at 95°C for 15 min and subjected to the described protocol.

### Protein extraction, SDS-PAGE and western blot

Proteins were extracted from 2 OD_600_ units of yeast cells as described in (Graf et al., 2017). Protein extracts were loaded on precast Mini-PROTEAN TGX precast gels (Bio-Rad). Proteins were blotted on a nitrocellulose membrane with the Trans-Blot Turbo Transfer System (Bio-Rad). The membrane was fixed with Ponceau S solution (P7170, Sigma Aldrich) and blocked for 1 h with 5% skim milk in 1xPBS containing 0.001% Tween-20 (PBS-T). The primary antibodies were incubated overnight in 5% skim milk in PBS-T. Peroxidase coupled secondary antibodies were incubated for 1h at room temperature. Antibodies are listed in ***Table S4***. The western blots were developed using the Super Signal West Pico Chemiluminescent Substrate (Thermo Scientific) and the ChemiDoc™ Touch Imaging System (Bio-Rad).

### Construction of strains with auxin-inducible degron

Strains carrying the auxin-inducible degron (AID*) for RNase H2 (catalytic subunit Rnh201) were created as described before (Morawska & Ulrich, 2013). Plasmids and oligonucleotides are listed in ***Table S3***.

### Construction of the cell cycle restricted RNase H2 alleles

The *S-RNH202-TAP-HIS3* and *G2-RNH202-TAP-HIS3* alleles were described previously (Lockhart et al., 2019).

The *G1-RNH202-TAP-HIS3* allele was created by amplifying the “G1 cassette” using the oligos oNA21 and oNA22 with the template pBL603 (containing the *SIC1* promoter, the first 315 bp of the *SIC1* gene and the NAT resistance cassette) (Johnson et al., 2016) by PCR (Janke et al., 2004). Transformed colonies were grown under selective pressure and sequence verified by sequencing with the respective oligonucleotide pairs. The cell cycle specific expression was confirmed by western blot. Plasmids and oligonucleotides are listed in ***Table S3***.

### α-factor arrest and release

For cell cycle analysis, cells were synchronized in G1 phase by addition of 4 µg/mL α-factor (Zymo research, mating hormone peptide) for 2 h. Cells were then spun and washed three times with water, released into fresh YPD medium and further grown at 25°C in a water bath. Protein and Flow cytometry samples were collected at indicated time points.

### Canavanine mutagenesis assay

The *CAN1* fluctuation analysis was performed as described in (Marsischky et al., 1996). Relevant genotypes for the *Can^R^* mutation assay were streaked out 48 h prior inoculation to conserve population doublings within replicates. At least 14 independent single colonies from each genotype were entirely excised from the agar plate using a sterile scalpel to inoculate a 10 ml of YPD medium. The cultures were incubated at 30°C, 250 rpm for 16h. After measuring the optical density of the cultures they were harvested by centrifugation. Then, each culture was resuspended in 1 ml sterile water. Exactly 1 ml of each resuspension was transferred to a new tube. From this, a 10-fold dilution series up to a dilution factor of 10^6^ was performed in a 96-well plate. Finally, 100 µl of all strains from the 10^-6^ dilution were plated on a YPD plate and distributed with exactly four glass beads per plate. All strains were plated on SC-ARG plates supplemented with 60 µg/mL canavanine with the indicated dilution factor. The plates were incubated for 72 h at 30°C before the outgrown colonies were manually counted. The medium for the *Can^R^* mutation assay was mixed, autoclaved and poured each day before plating to maintain constant conditions between replicates.

For evaluation, the number (#) of mutant cells per culture, representing *r*, was calculated:

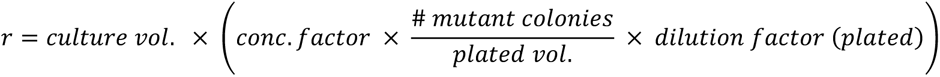

The following correction was used to account for the progenies of each individual *CAN1* mutation event per cell. With *M* being a scaled value that represents the number of cells that have actually undergone a mutation event (from which the counted progenies originated):

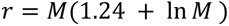

The final mutation rate was calculated dividing *M* by the total number of cells present in the initial culture:

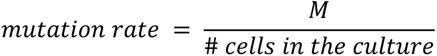

The data was plotted as the Median with 95% Confidence interval using the GraphPad PRISM8 software.

### Plating Assay

Exponential cultures at 30°C were synchronized with α-factor for 1h and then split to start the degradation of Rnh201-AID*-9Myc with 1 mM auxin in 50% of the samples during the residual 1h of synchronization. Half of the culture remained arrested in G1 phase and the other half was released into S phase, by washing out α-factor, in the presence or absence of 1 mM auxin for 30 min. Of each culture and condition, a suitable dilution was empirically determined that yielded in 100-200 colonies per YPD agar plate after outgrowth. The plates were incubated for 2 days at 30°C. Colonies of 7 replicates were manually counted and adjusted for differences in optical density (OD_600_) before dilution. Statistical analysis and plot generation was performed using Prism8 (GraphPad Software).

### Yeast spot assay

Single colony derived yeast cells were incubated overnight at the appropriate temperature in liquid medium. Cells were diluted to 0.5 OD_600_ and spotted in ten-fold serial dilutions onto YPD plates, SC plates, or plates containing the indicated amount of genotoxic drugs, i.e. methyl methane sulfonate (MMS), Camptothecin (CPT) or hydroxyurea (HU) (all drugs: Sigma-Aldrich). The agar plates were incubated at the indicated temperatures and time and imaged using the ChemiDoc™ Touch Imaging System (Bio-Rad).

Standard YPD agar has pH 5.5. To make YPD agar plates with alkaline pH, we titrated melted YPD agar with 10N NaOH until pH 8.0 was reached. The *wsc1::KAN* knockout strain was used as a positive control for the alkaline agar plates (Serra-Cardona et al., 2015).

### Alkaline Gel electrophoresis

Analysis of alkaline-labile sites in genomic DNA was performed as reported earlier (Nick McElhinny et al., 2010).

### Construction of cell cycle regulated *RNH202* allele in the SGA query strain background

The *G1-RNH202-TAP (this study)*, *S-*, and *G2-RNH202-TAP* alleles (Lockhart et al., 2019) were crossed to the haploid background strain (Y8205, Source C. Boone) for the SGA query strain construction. Selection of diploids, sporulation and tetrad analysis generated the four query strains used in SGA analysis. Cell cycle restricted protein expression of Rnh202 was confirmed by arrest and release experiment and western blot analysis. The selectable markers for SGA analysis were verified by PCR (oMT86/oMT91 for *can1Δ::STE2pr-Sp_his5*, oMT89/oMT90 for *lyp1Δ::STE3pr-LEU2*) and replica-plating on YPD + (50 μg/ml canavanine, 50 μg/ml thialysine). The yeast strains are listed in ***Table S2***.

### Synthetic genetic array (SGA) screen procedure and data evaluation

The *G1-*, *S-*, and *G2-RNH202-TAP* query strains and a wild type *RNH202* control query were crossed with the haploid genome-wide library of yeast gene deletion mutants, the YKO (Winzeler et al., 1999). Crosses were performed in 1536-colony format, with the four queries combined on each screen plate, with four technical replicates of each cross, arranged next to each other. To minimize spatial effects, four outer rows and columns contained dummy strains. Mating, sporulation and selection of haploids carrying both a query allele (cell cycle regulated *RNH202* alleles or wild type control) and a gene deletion were performed by sequential pinning of yeast colonies on appropriate selective media using a RoToR pinning robot (Singer Instruments) as described (Baryshnikova et al., 2010). Plates with the final colony arrays were imaged after 24 h with the Singer PhenoBooth colony imager. Data analysis was performed in R (R Core Team (2021). R: A language and environment for statistical computing. http://www.R-project.org/) as detailed in the R vignette (***S6 HMTL***). Briefly, photographs of colony arrays were segmented using the gitter package (Wagih & Parts, 2014) to determine colony size. Measurements of empty positions and four outer rows and columns were assigned NA values. Colony size measurements on each plate were corrected for spatial effects using the SGA tools package (Wagih & Parts, 2014) and normalized to the median on each plate. Genetic interactions in double mutants were identified under the assumption of multiplicative combination of effects of single mutants in the absence of genetic interactions (Baryshnikova et al., 2010). For that, normalized colony size measurements were divided by the median per query to obtain normalized double mutant fitness. For each mutant in the YKO collection, differences between crosses with a cell cycle and the wild type queries were assessed with a t-test, excluding replicates contributing more than 90% of variance. The p-values were adjusted for multiple testing using the Benjamini-Hochberg method. Finally, replicates were summarized by their mean, excluding replicates contributing more than 90% of variance (***S7 XLS***). Negative genetic interactions were verified through manual generation of haploid double mutants, by crossing single colonies from the YKO haploid collection to the *S-RNH202* allele, selection for diploids, sporulation and tetrad analysis. False positives and linked genes have been excluded in the final analysis (greyed out).

### Materials

Materials such as antibodies, enzymes, and chemicals are listed in ***Table S4***.

### Numerical data

Underlying numerical data for all graphs and source data for western blots are provided in ***Table S5***.

## Author contributions

Conceptualization: B.L., V.K., N.S.

Alkaline gels, plating assay, SGA screen: M.T.

SGA screen evaluation: M.T., J.J.F., A.K.

Spottings: V.K., M.T., N.S.

Tetrad dissections: M.T., N.S., A.L.

Flow cytometry: M.T., N.S., S.M.

Helped design the research: T.J., P.B., O.V.

Writing of the original draft: N.S., B.L.; all authors read and approved the final version of the manuscript.

Supervision: B.L., P.B., A.K.

Funding acquisition: B.L.

## Acknowledgements

We thank the Luke lab members for support and discussions and the Media lab, Flow cytometry and Protein Production core facilities of the Institute of Molecular Biology (IMB gGmbH) for technical support. We thank Stefanie Reimann, Simone Snead, Rebecca Medina, Lara Perez-Martinez, Despina Giamaki, Nina Lohner, and Dennis Knorr for technical assistance. We thank the Ulrich lab at IMB for reagent exchange and sharing equipment. Thanks to H. Hombauer (DKFZ Heidelberg) for the help with establishing the Canavanine mutagenesis assays. Thanks to Georg Stoecklin (Heidelberg University) for stimulating the alkaline agar plate pH assay. We thank the Zhang lab for sharing histone mutant plasmids (Han et al., 2013).The model illustration was created with BioRender.com. B.L. receives funding from the Heisenberg Program of the DFG - LU 1709/2-1 (B.L.) and from the Deutsche Forschungsgemeinschaft (DFG, German Research Foundation) – Project-ID 393547839 – SFB 1361.

## Abbreviations

rNMP: ribonucleoside monophosphate
RER: ribonucleotide excision repair
seDSB: single-ended double-strand breaks
NLR: rNMP-derived nick lesion repair
HDR: homology-directed repair
gDNA: genomic DNA
ssDNA: single-strand DNA
ssDNA break: nick
Top1: Topoisomerase 1
R-loop: RNA-DNA hybrid with displaced ssDNA strand
CPT: Camptothecin, Top1 poison, forms Top1-DNA-covalent complexes
HU: hydroxyurea, ribonucleotide reductase inhibitor
MMS: methyl methane sulfonate, alkylation agent
AID*: auxin-inducible degron
IAA: indole acetic acid, auxin
CRL: Cullin-RING ubiquitin ligase
CRL4: Cullin-4 (human homolog of Rtt101)
RCNA: replisome coupled nucleosome assembly
SL: synthetic lethal

## Supporting information

**Figure S1.**
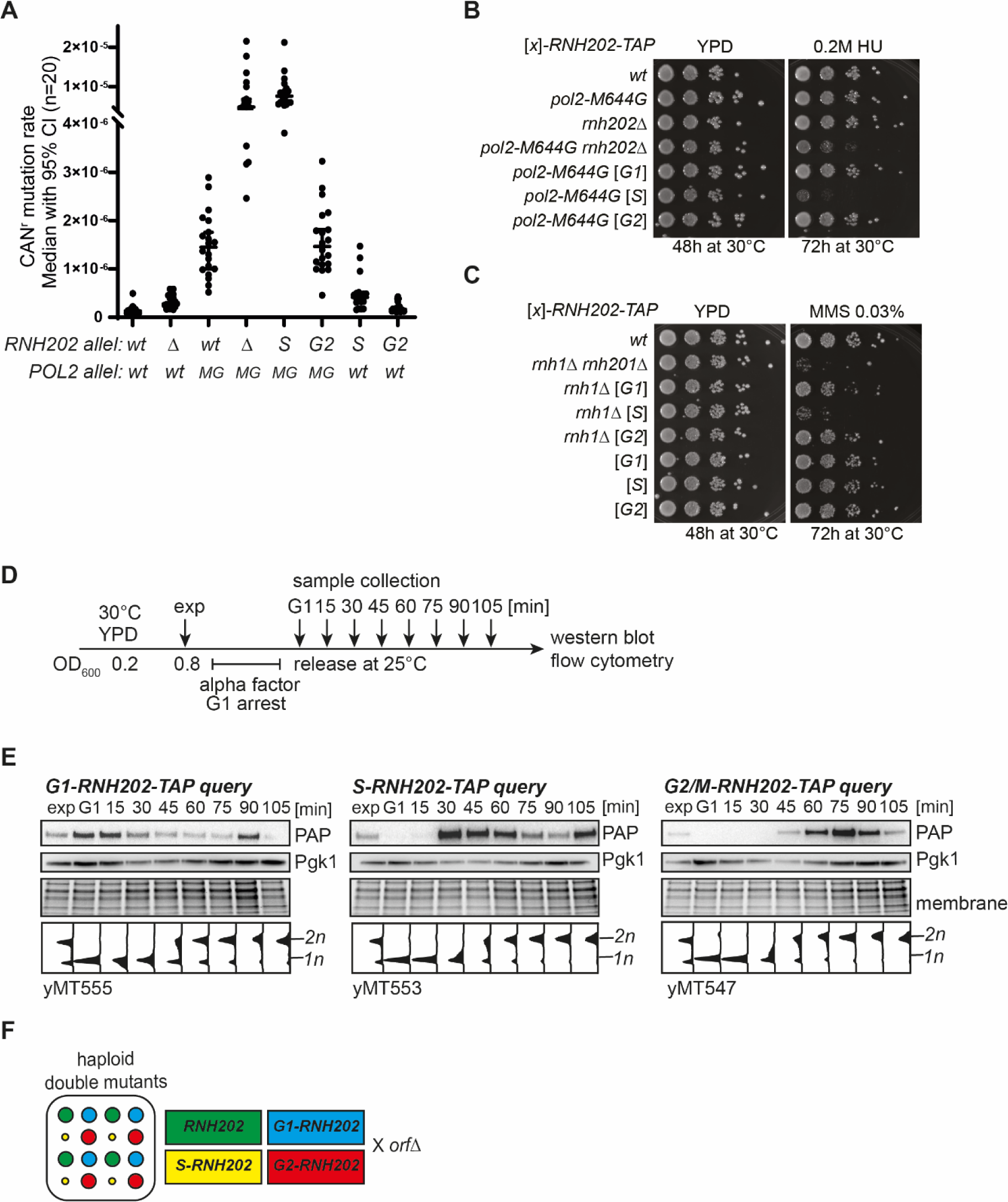
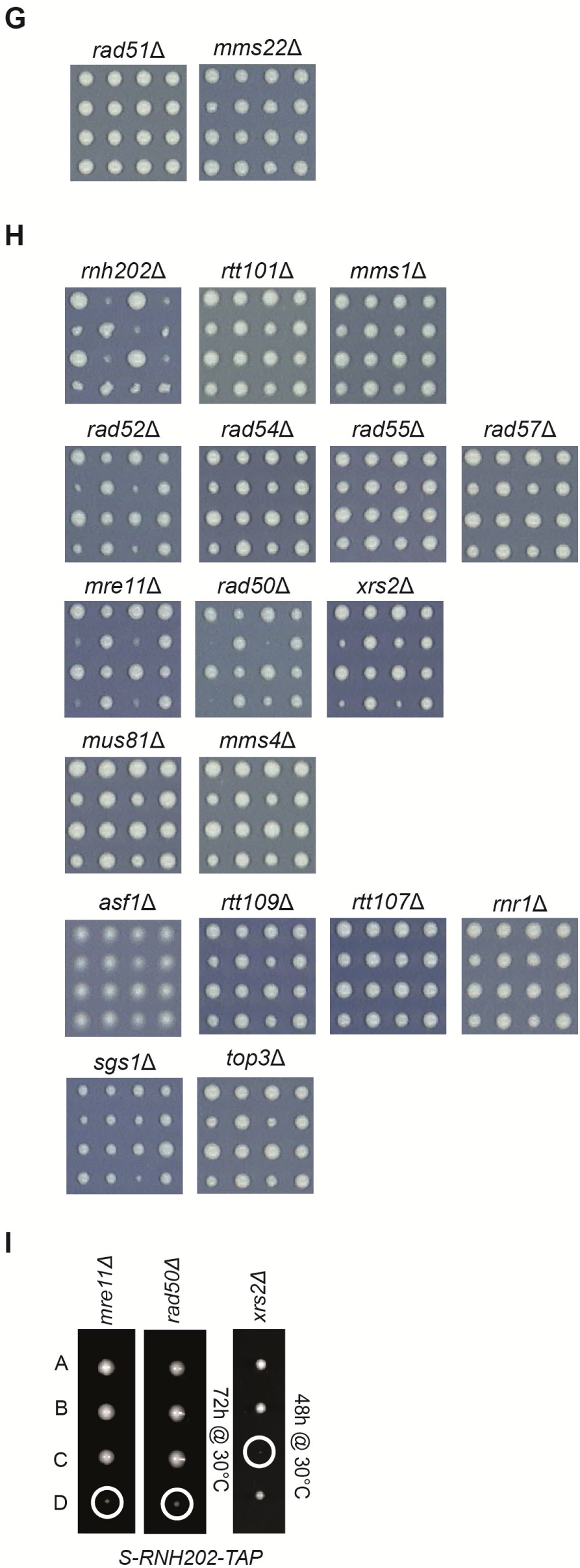
(A-C) Features of the *S-RNH202* allele. **(A)** Fluctuation assay to measure the Canavanine-resistance mutagenesis comparing the indicated genotypes. The assay showed that the *S-RNH202-TAP* allele has the same mutagenesis load as the RER-deficient *rnh202*Δ strain in the presence of high rNMP load induced by the *pol2-M644G* allele. (**B)** Tenfold serial dilution of the indicated mutants to compare the cell cycle alleles of *RNH202-TAP* with the deletion *rnh202*Δ in the presence of high rNMP load (*pol2-M644G*) to compare the rNMP-dependent toxicity. Spontaneous hydrolysis of persistent rNMPs in the RER-deficient *pol2-M644G rnh202*Δ led to reduced viability, while the active rNMP-nicking in the *pol2-M644G S-RNH202-TAP* mutant was inviable in the presence of hydroxyurea (HU). Note that *G1-RNH202-TAP* and *G2-RNH202-TAP* alleles support full viability. **(C)** Serial dilution of the indicated *RNH202-TAP* cell cycle alleles in combination with *RNH1* knockout to assess the R-loop removal activity of the *RNH202-TAP* cell cycle alleles in the presence of MMS, a drug that accumulates R-loop levels in cells. **(D-H) Query strain characterization and representative SGA raw data. (D)** To test the cell cycle specific expression of the query strains generated for the SGA screen, cells were arrested in the G1 phase with α-factor at 30°C. Upon full synchronization, the culture was released into the cell cycle at 25°C and samples for western blot and DNA content analyses were collected every 15 min. **(E)** The *RNH202* query alleles are tagged with tandem affinity purification (TAP) tags that allow the detection of protein expression using a PAP antibody. The Pgk1 antibody and the Ponceau Red stained nitrocellulose membrane served as loading controls. The strains used here have the mating type *MATa* (susceptible to α-factor arrest); for the SGA screen in Figure 1, the corresponding *MATalpha* strains from the same dissection were used. **(F)** Schematic overview of genotypes in a SGA panel. Every set was pinned in four replicates. **(G)** Selected example panels from the raw SGA screen data that scored below threshold, but were manually verified due to their context with the candidate genes in Figure 1G. **(H)** Selected example panels from the raw SGA screen data that scored above threshold and were manually verified. **(I)** Manual tetrad dissection confirmation of the requirement for the components of the MRX complex for *S-RNH202* survival as the *mre11*Δ *S-RNH202*, *rad50*Δ *S-RNH202*, and the *xrs2*Δ *S-RNH202* double mutants are inviable.

**Figure S2.**
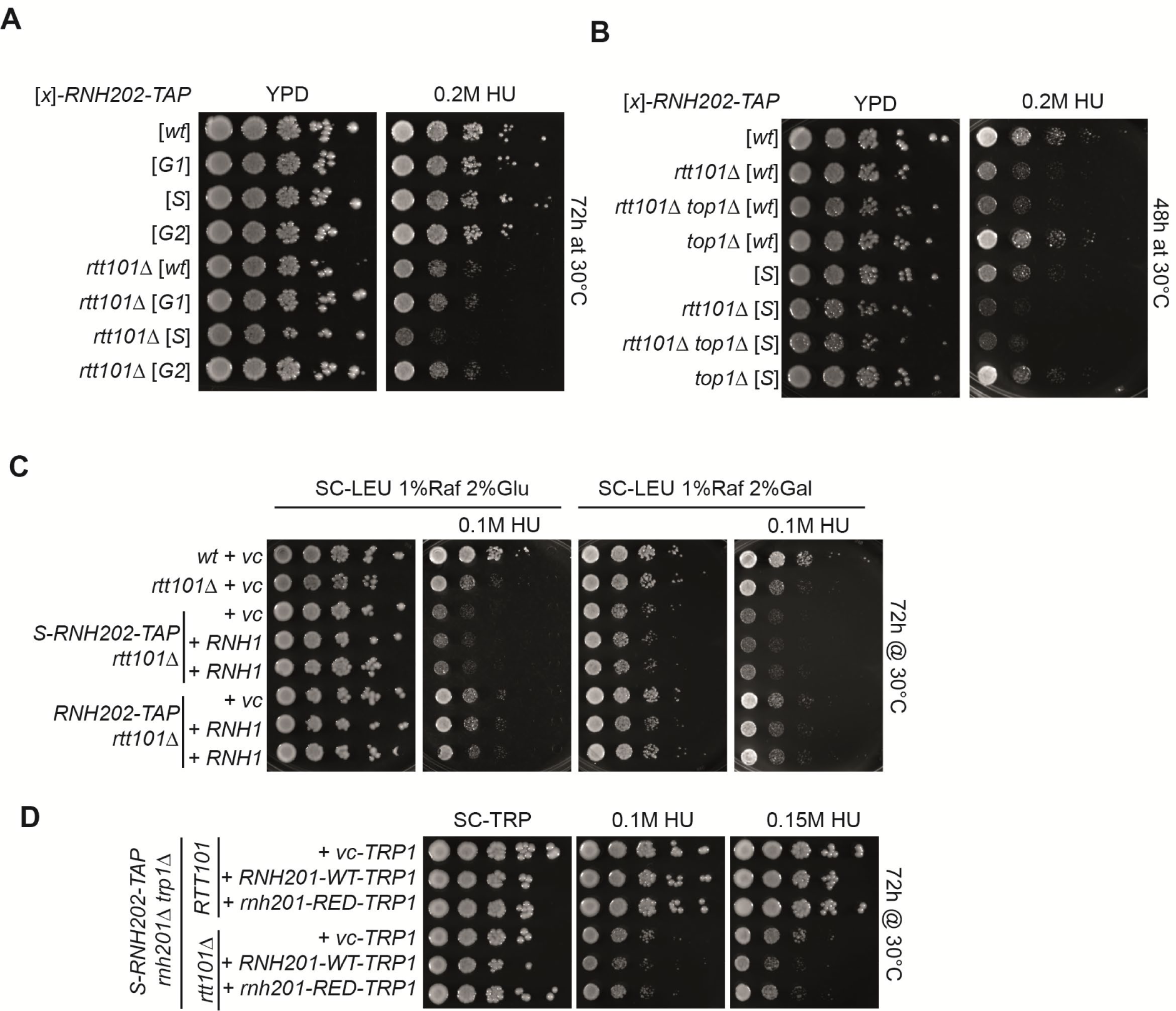
Loss of *RTT101* causes Top1-independent drop in viability in conditions of increased genomic rNMP hydrolysis. **(A)** The *rtt101*Δ *G1-RNH202* and *rtt101*Δ *G2-RNH202* double mutants are epistatic in terms of HU sensitivity compared to *rtt101*Δ alone. The *S-RNH202 rtt101*Δ double mutant is sick in the presence of HU. After 48h the *S-RNH202-TAP rtt101*Δ double mutant growth is delayed which is not visible after 72h incubation. However, in the presence of HU, the S-*RNH202-TAP rtt101*Δ double mutant strain was very sick. **(B)** Deletion of the *TOP1* gene does not rescue of the toxicity of *S-RNH202-TAP rtt101*Δ in the presence of HU, showing that this is a Top1 independent toxicity. Images were taken after 2 days of growth at 30°C. **(C)** Spot assay using *RNH1* overexpression to test the effect of R-loop removal in the *S-RNH202-TAP rtt101*Δ double mutant. **(D)** Spot assay to demonstrate that the rNMP-excision function of S phase restricted *RNH202* is causing the toxicity in the *S-RNH202-TAP rtt101*Δ double mutant. We generated the *S-RNH202-TAP rnh201*Δ *rtt101*Δ *trp1*Δ quadruple mutant that was transformed with *pRS416-RNH201-WT-URA3*. Subsequently, the strains were co-transformed with *pRS413-vc-TRP1, pRS413-RNH201-WT-TRP1*, or *pRS413-rnh201-RED* plasmids. Then, we selected against for loss of the *pRS416-RNH201-WT-URA3* plasmid in the presence of 5-FOA and spotted the resulting strains. Images were taken after 2 or 3 days of growth at 30°C. HU = hydroxyurea

**Figure S3.**
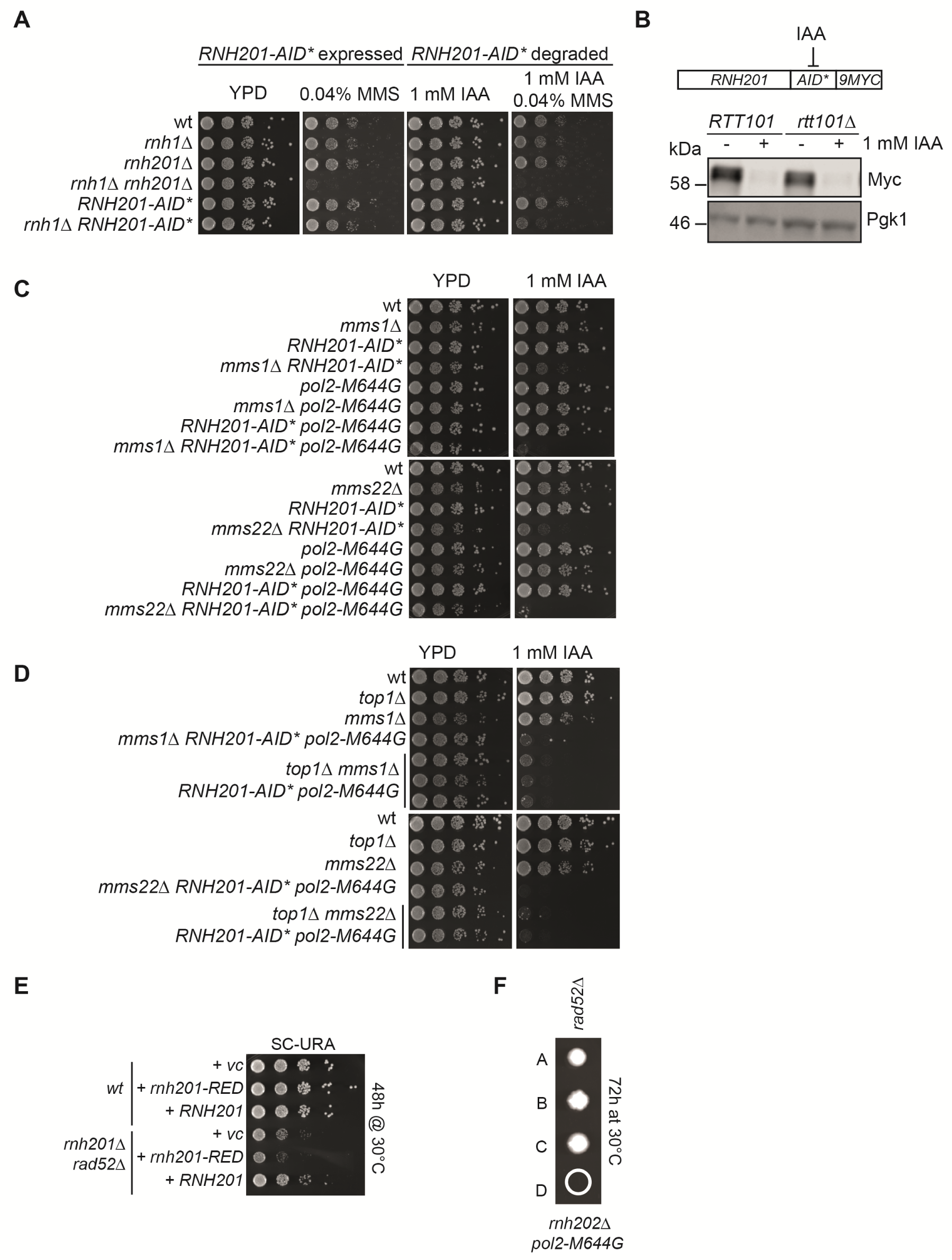
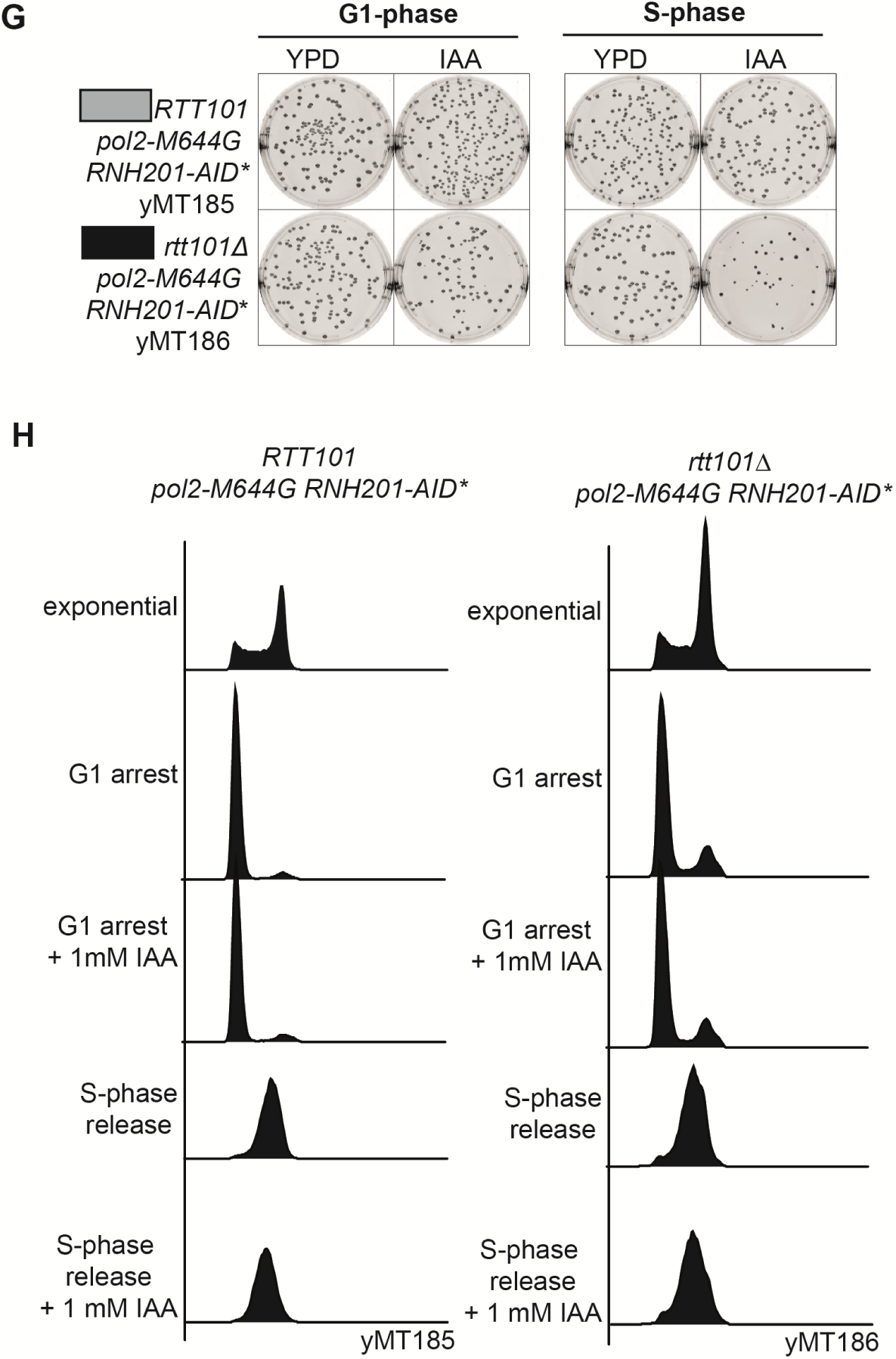
The synthetic sickness of the Rtt101^Mms1-Mms22^ complex with RNase H2-deficiency is Top1-independent. **(A)** Functionality test of the *RNH201-AID\** strain by serial dilution spot assay. The *rnh1*Δ *rnh201*Δ double mutant is sensitive to MMS (Lazzaro et al., 2012). The depletion of *RNH201-AID\** by auxin in the *rnh1*Δ background impaired cell growth. **(B)** The auxin-inducible degron (AID*) tag results in proteasomal degradation of the fusion protein in the presence of auxin (Morawska & Ulrich, 2013). The western blot of exponential cells treated for 1 h with 1 mM IAA confirmed a robust degradation of the Rnh201-AID*-9Myc protein. **(C)** The depletion of *RNH201-AID\** in the presence of auxin resulted in a synthetic sick growth phenotype with *mms1*Δ, and *mms1*Δ, which was amplified into a synthetic lethal phenotype when combined with high genomic rNMP load (*pol2-M644G*). **(D)** The synthetic lethality of the *pol2-M644G mms1*Δ *RNH201-AID\** and the *pol2-M644G mms22 RNH201-AID\**triple mutant on auxin plates was Top1-independent. Images were taken after 2 days of growth at 30°C. MMS = methyl methane sulfonate, IAA = indole acetic acid (auxin). **(E)** Spot assay showing the effect of *RAD52* deletion in the RER-deficient RNase H2 mutant (*rnh201-RED*). **(F)** Tetrad dissection showed that *rnh202*Δ *pol2-M644G* double mutants require *RAD52* for survival. **(G-H) Colony formation assay representative images and DNA profiles. (G)** Images of representative agar plates from the colony formation assay. Summary of the data shown in Figure 3 F-H. **(H)** The DNA profiles of the strains used for the colony formation assay in (G) and Figure 3 F-H were measured by flow cytometry with Sytox Green stained cells.

**Table S1**

SGA screen hit (Figure 1) evaluation by manual dissection. Representative tetrad dissection images in Figure S1. The exclusive G2-RNH202 allele hits, *YLR236* and *FYV10* were not verified but greyed out in the string network in Figure 1G as they were not genetically interacting with the *S-RNH202* allele.

**Table S2**

Yeast strains used in this study. (xlsx)

**Table S3**

Plasmids and oligonucleotides used in this study. (xlsx)

**Table S4**

Materials used in this study. (xlsx)

**Table S5**

Underlying numerical data. (xlsx)

**S6 Html**

R vignette for SGA analysis

**S7 Xls**

*RNH202*_cell cycle KO screen data

